# Reducing batch effects in single cell chromatin accessibility measurements by pooled transposition with MULTI-ATAC

**DOI:** 10.1101/2025.02.14.638353

**Authors:** Daniel N. Conrad, Kiet T. Phong, Ekaterina Korotkevich, Christopher S. McGinnis, Qin Zhu, Eric D. Chow, Zev J. Gartner

**Affiliations:** Department of Pharmaceutical Chemistry, University of California, San Francisco, San Francisco, CA 94158, USA; Department of Biochemistry and Biophysics, University of California, San Francisco, San Francisco, CA 94158, USA; Department of Pathology, Stanford University, Stanford, CA 94305, USA; Gladstone-UCSF Institute of Genomic Immunology, San Francisco, CA 94158, USA; Parker Institute for Cancer Immunotherapy, San Francisco, CA 94129, USA; Laboratory for Genomics Research, San Francisco, CA, 94158, USA; Chan Zuckerberg Biohub, San Francisco, CA 94158, USA; Center for Cellular Construction, University of California, San Francisco, CA 94158, USA

## Abstract

Large-scale scATAC-seq experiments are challenging because of their costs, lengthy protocols, and confounding batch effects. Several sample multiplexing technologies aim to address these challenges, but do not remove batch effects introduced when performing transposition reactions in parallel. We demonstrate that sample-to-sample variability in nuclei-to-Tn5 ratios is a major cause of batch effects and develop MULTI-ATAC, a multiplexing method that pools samples prior to transposition, as a solution. MULTI-ATAC provides high accuracy in sample classification and doublet detection while eliminating batch effects associated with variable nucleus-to-Tn5 ratio. We illustrate the power of MULTI-ATAC by performing a 96-plex multiomic drug assay targeting epigenetic remodelers in a model of primary immune cell activation, uncovering tens of thousands of drug-responsive chromatin regions, cell-type specific effects, and potent differences between matched inhibitors and degraders. MULTI-ATAC therefore enables batch-free and scalable scATAC-seq workflows, providing deeper insights into complex biological processes and potential therapeutic targets.

## Introduction

Single-cell genomics techniques allow for the composition and state of complex systems to be compared across time, space, individual, and perturbation. Fundamental challenges of these methods include the high reagent costs, time, and technical artifacts (e.g. batch effects) associated with their complex workflows. Sample multiplexing technologies circumvent these challenges, reducing the complexity of experiments and eliminating batch effects by pooling samples and processing them together through downstream molecular biology steps. Such methods are now widely used to generate high throughput single-cell RNA-seq (scRNA-seq) datasets and enable transcriptomic profiling of dozens to hundreds of samples at once^1–4^. Analogous multiplexing methods have also been described recently for single-cell assay for transposase-accessible chromatin (scATAC-seq)^5–11^, an epigenomic analysis technique that measures regions of open chromatin in individual cells using Tn5 transposase loaded with sequencing adapters. Notably, the majority of existing scATAC-seq sample multiplexing methods require each sample to be transposed independently or even split across many individual reactions, limiting assay scalability and increasing experiment costs.

Beyond these limitations of parallel transposition workflows, variability in the nuclei:Tn5 ratio between samples can introduce significant batch effects that confound downstream analysis (Fig. 1A). Tn5 is a single-turnover enzyme, so the stoichiometric ratio of Tn5 to nuclei dictates the average number of fragments generated per nucleus in a reaction; this can even bias the proportions of genomic features detected^12–14^. While this phenomenon is well-established in bulk ATAC-seq workflows, how variable nuclei:Tn5 ratios contribute to batch effects in scATAC-seq analysis has not been thoroughly explored.

**Figure 1.**
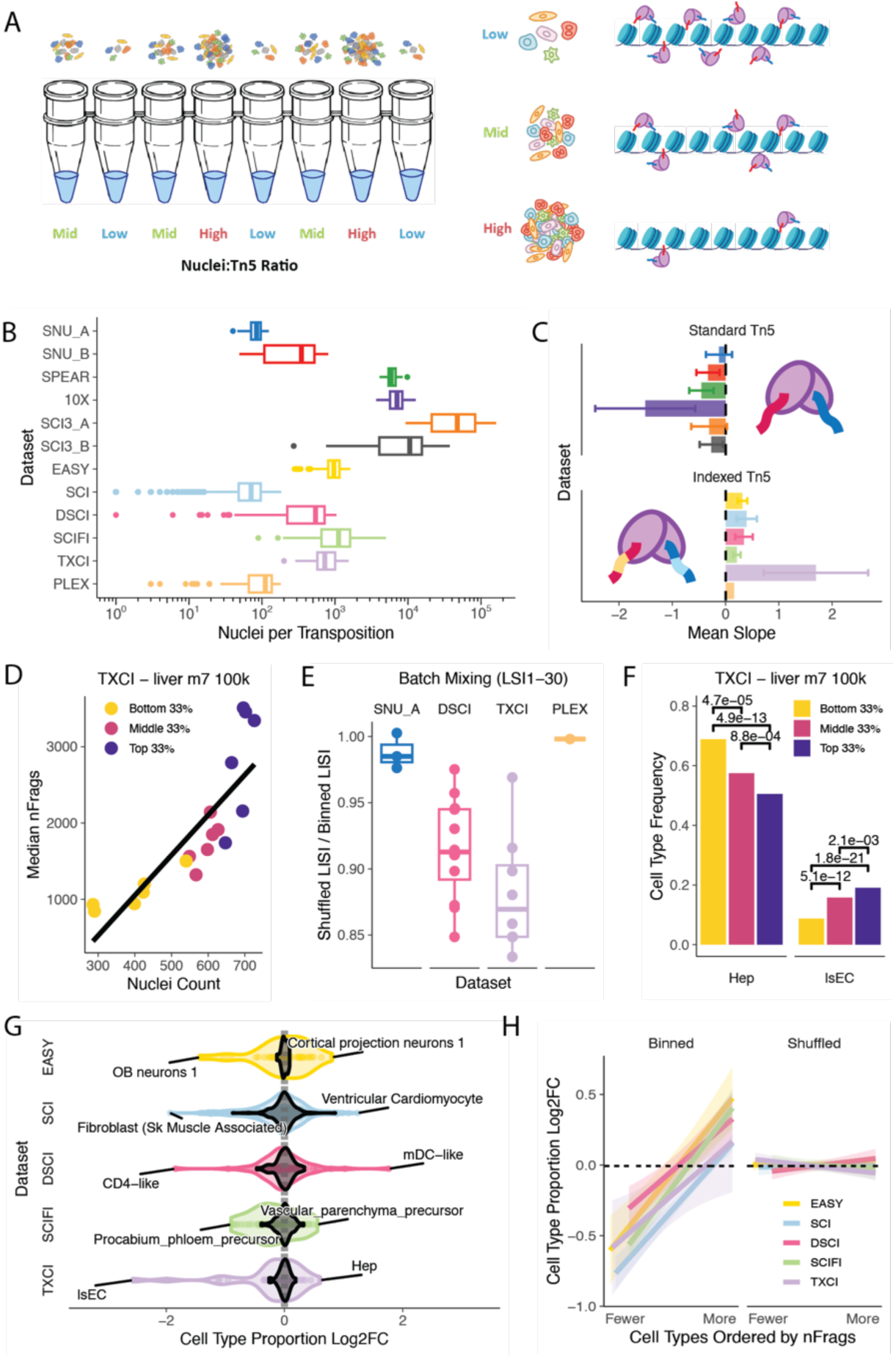
Transposition batch size effects in published datasets. A. Variable nuclei counts in separately-transposed samples bias the number of cuts made per nucleus, determining per-nucleus fragment yield B. Inspection of 12 published datasets shows considerable variation in transposition batch size within individual experiments and datasets C. Methods using standard Tn5 (non-indexed adapter oligos) exhibit a negative association between transposition batch size and median per-nucleus fragment count, while methods using indexed Tn5 exhibit an unexpected positive association. D. Example sample from the TXCI dataset. Points represent the nuclei count and median fragment count per transposition reaction, and are colored by transposition batch size tercile. E. Relative mixing of transposition batch size terciles in the 30-dimensional LSI reduction across 4 datasets. Points represent separate biological samples and/or technical replicates per dataset. Average Local Inverse Simpon’s Index (LISI) scores per sample were normalized to “idealized” mixing scores derived by permuting tercile labels. F. Two demonstrative cell types from the sample in D), showing statistically significant changes in cell type frequency according to transposition batch tercile. P-values represent results from two-sided Chi-squared proportion tests. G. Log2 fold-changes in cell type proportions between the bottom and top transposition batch size terciles plotted for all prominent cell types (> 5% of sample) across all samples of 5 datasets. For comparison, Log2 fold-changes were computed after permuting tercile labels (black). H. The same log2 fold-changes reported in G), plotted as a function of increasing mean fragment yield for each individual cell

Here, we describe MULTI-ATAC, a scATAC-seq sample multiplexing technology that improves scATAC-seq sample throughput and optimizes scATAC-seq data quality through doublet detection and the mitigation of batch effects caused by variable nuclei:Tn5 ratios. First, we re-analyzed publicly-available scATAC-seq datasets and identified the presence of significant batch effects that arise due to variable nuclei:Tn5 ratios. Second, we demonstrate that MULTI-ATAC is compatible with pooled transposition workflows and enables the generation of multiplexed scATAC-seq data with minimal batch effects. Finally, we leverage MULTI-ATAC to perform a 96-plex multiomic drug perturbation experiment measuring how primary human immune cells respond to diverse inhibitors and proteolysis targeting chimeras (PROTACs) targeting chromatin remodeling enzymes. From these data we identify tens of thousands of immune- and drug-responsive chromatin regions and genes and discover that MS177 accentuates NF-κB signaling, while SWI/SNF perturbation induces a potent type I interferon response.

## Results

### Transposition batch effects detected in published datasets

To determine if batch effects are linked to nuclei:Tn5 ratio in large-scale and multi-sample scATAC-seq experiments, we re-analyzed 12 publicly-available datasets representing a variety of species and library preparation methods (Table 1) and assessed the magnitude of batch effects between independent transposition reactions in each dataset^5,7,8,10,11,15–20^ (Methods). Importantly, we made the assumption that the number of nuclei in the dataset associated with each Tn5 reaction was correlated to the number of nuclei used as input. The range of nuclei per sample varied greatly within a single experiment, spanning a range of 2-fold to 66-fold (Fig. 1B; Table 1), and thereby offered the opportunity to quantitatively measure batch effects between samples. Notably, datasets generated from experiments where low numbers of samples were split across many transposition reactions – a situation where nuclei counts are easiest to control – had minimal nuclei count variability. Conversely, datasets from experiments with high numbers of unique samples or where nuclei were isolated from tissue samples – a situation where nuclei counts are challenging to control – had far greater nuclei count variability between transposition reactions. These observations across 12 datasets suggest that nuclei count variability in transposition reactions is an intrinsic feature of complex scATAC-seq experiments.

**Table 1.**
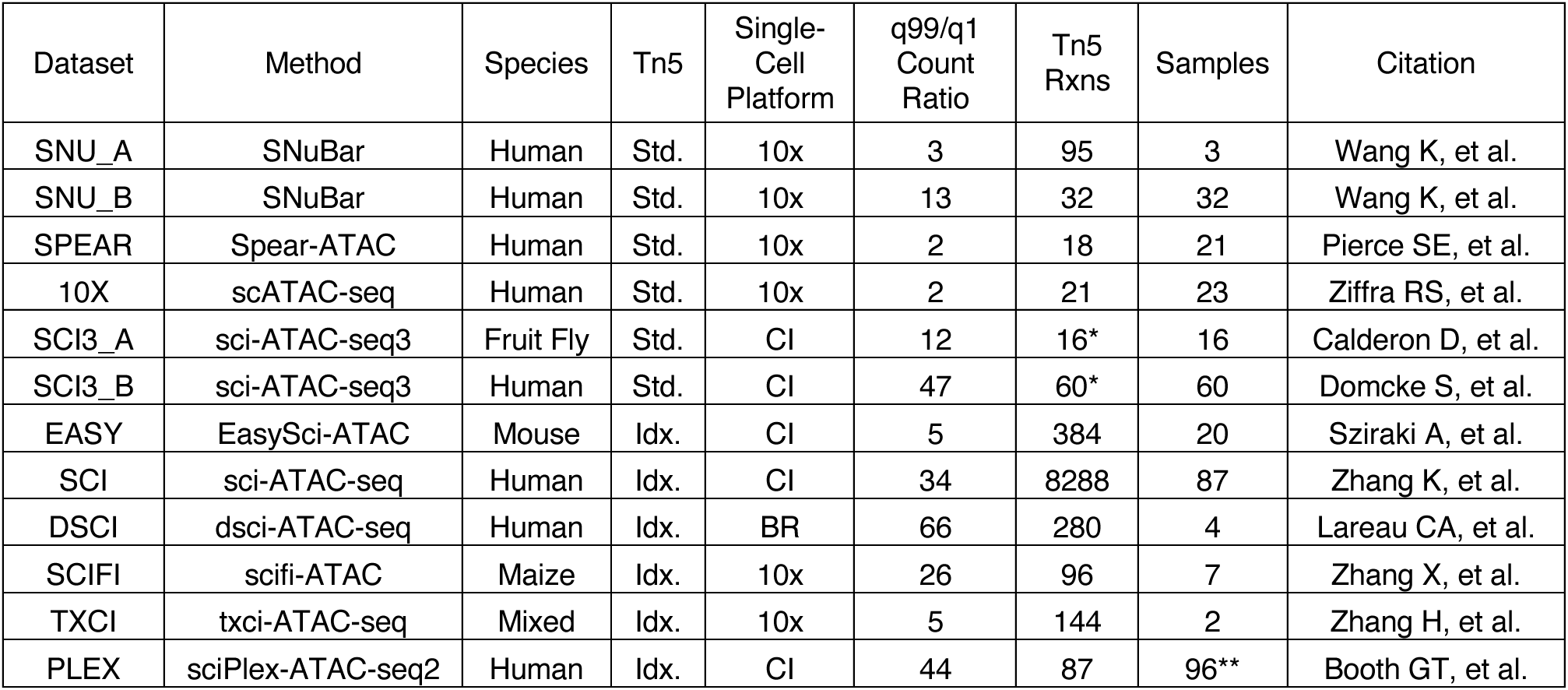
Published datasets reanalyzed for transposition batch effects. Single-cell ATAC-seq datasets from 11 publications spanning a variety of different techniques and biological systems. The number of nuclei per transposition reaction in each dataset was tabulated, and the range of transposition batch sizes was represented by the ratio of the maximum and minimum nuclei counts (excluding outliers above and below the 99^th^ and 1^st^ quantile of the count distribution, respectively). The number of transposition reactions represents the total recovered in the final dataset, and at times is less than the original experimental design intended due to drop-outs. * sci-ATAC-seq3 datasets (SCI3_A and SCI3_B) actually reflect aggregations of 11 and 4 transposition reactions per sample, respectively, due to sci-ATAC-seq3 methodology ** PLEX reflects 96 samples pooled and split across 96 individual reactions

| Dataset | Method | Species | Tn5 | Single-Cell Platform | q99/q1 Count Ratio | Tn5 Rxns | Samples | Citation |
| --- | --- | --- | --- | --- | --- | --- | --- | --- |
| SNU_A | SNuBar | Human | Std. | 10x | 3 | 95 | 3 | Wang K, et al. |
| SNU_B | SNuBar | Human | Std. | 10x | 13 | 32 | 32 | Wang K, et al. |
| SPEAR | Spear-ATAC | Human | Std. | 10x | 2 | 18 | 21 | Pierce SE, et al. |
| 10X | scATAC-seq | Human | Std. | 10x | 2 | 21 | 23 | Ziffra RS, et al. |
| SCI3_A | sci-ATAC-seq3 | Fruit Fly | Std. | CI | 12 | 16* | 16 | Calderon D, et al. |
| SCI3_B | sci-ATAC-seq3 | Human | Std. | CI | 47 | 60* | 60 | Domcke S, et al. |
| EASY | EasySci-ATAC | Mouse | Idx. | CI | 5 | 384 | 20 | Sziraki A, et al. |
| SCI | sci-ATAC-seq | Human | Idx. | CI | 34 | 8288 | 87 | Zhang K, et al. |
| DSCI | dsci-ATAC-seq | Human | Idx. | BR | 66 | 280 | 4 | Lareau CA, et al. |
| SCIFI | scifi-ATAC | Maize | Idx. | 10x | 26 | 96 | 7 | Zhang X, et al. |
| TXCI | txci-ATAC-seq | Mixed | Idx. | 10x | 5 | 144 | 2 | Zhang H, et al. |
| PLEX | sciPlex-ATAC-seq2 | Human | Idx. | CI | 44 | 87 | 96** | Booth GT, et al. |

We next asked whether data quality-control metrics correlated with the number of nuclei processed per reaction. scATAC-seq methods can be divided into two classes depending on whether they utilize Tn5 loaded with barcoded adapters (‘indexed transposome’) or universal adapters (‘standard transposome’). In standard transposome datasets, we observed that the median number of fragments per cell was negatively correlated with the number of transposed nuclei (Fig. 1C, Fig. S1A), mirroring results in bulk ATAC-seq^12^. Interestingly, indexed transposome datasets exhibited the opposite trend, yielding more fragments per cell in batches with greater nuclei counts (Fig. 1C, Fig. S1A). While the mechanism underlying this trend reversal remains unclear, ‘index hopping’ between transposition products due to the presence of free adapters could play a role^10,11^.

Regardless of the mechanism or direction of the relationship, a correlation between transposition batch size and fragment yield could be detrimental to analysis as previously described in bulk ATAC-seq data. We therefore investigated how this technical artifact impacted downstream analyses and biological interpretation. Dimensionality reduction is commonly used during scATAC-seq analysis and provides the foundation for unsupervised clustering, cell type annotation, and differential accessibility analysis. Due to the inherent sparsity of chromatin accessibility data, Latent Semantic Indexing (LSI) is the predominant algorithm applied to scATAC-seq data^21,22^. In practice, the first LSI component correlates strongly with per-cell fragment counts, and is thus customarily excluded to avoid technical bias^10,18,21–24^. However, by separating cells by subtype, we find that many more LSI components covaried in absolute magnitude with per-cell fragment counts, indicating that simply excluding the first LSI component is not sufficient to abrogate depth-related effects on clustering (Fig. S1D-E).

To better quantify the impact of variable Tn5 batch size (and thus variable nuclei:Tn5 ratio) on dimensionality reduction, we selected datasets where unique samples were transposed across many reactions and for which fragment data were readily available (SNU_A, DSCI, TXCI, and PLEX). We binned the nuclei of each dataset into terciles according to Tn5 batch size (Fig. 1D, Fig. S1A). We then used the Local Inverse Simpson’s Index algorithm^25^ (LISI) to score the degree of batch mixing of the terciles of each dataset across 30 LSI dimensions, and compared this value to the degree of mixing when bin assignments were permuted to represent perfect mixing (Fig. 1E). Two of the datasets, SNU_A and PLEX, seemed largely unaffected; these datasets also exhibited the weakest association with transposition batch size (Fig. 1C), likely due in part to experimental designs that facilitated consistent loading of transposition reactions. The two datasets with significantly impacted batch mixing, DSCI and TXCI, represent more complex experiments where nuclei from multiple heterogeneous primary samples (bone marrow mononuclear cells, human lung, mouse liver/lung) were isolated separately and transposed across many reactions – resulting in much stronger correlations between Tn5 batch size and fragment counts (Fig. 1C, Fig. S1A). This supports the notion that only simple experimental designs that allow for precise control of nuclei counts can control for batch effects. Furthermore, excluding the first LSI component from this analysis yielded similar results, further supporting that bias from variation in per-cell library complexity is not uniquely captured by and removed with the first LSI component (Fig. S1B).

In addition to influencing dimensionality reduction, we also observed significant shifts in cell type composition between Tn5 batches (Fig. 1F, Fig. S1C). Specifically, across 5 datasets representing heterogeneous samples split across many individual transposition reactions, we observed that the proportions of highly-prevalent cell types (i.e., > 5% of the total) such as hepatocytes and sinusoidal endothelial cells in the TXCI dataset, varied considerably between Tn5 batch terciles (Fig. 1F, Fig. S1C). Importantly, the observed variation far exceeds differences in cell type proportions computed after permuting bin labels (Fig. 1G). One possible explanation for this result derives from differences in fragment yields among different cell types, in turn resulting in differential sensitivity to quality control filtering for cells with naturally lower fragment counts. Indeed, comparing the mean fragment count per cell type and its change in proportion between Tn5 bins revealed that cells with fewer fragments are selected against in Tn5 batches that yield fewer fragments (Fig. 1H). Collectively, these results suggest that the nuclei:Tn5 ratio during transposition can dramatically influence two critical steps of scATAC-seq analysis and therefore biological interpretation.

### MULTI-ATAC barcoding accurately classifies sample-of-origin and doublets

A simple solution to avoid batch effects from variable nuclei:Tn5 transposition ratios would be a sample multiplexing strategy that enables all samples to be transposed in a single pool, additionally streamlining the workflow and minimizing reagent costs. In order for samples to be pooled during transposition, sample-specific DNA barcodes must be incorporated into or onto nuclei in a manner that survives the transposition incubation without interfering with the reaction itself. In pursuit of this goal, we adapted the previously described MULTI-seq^1^ barcoding strategy to be compatible with scATAC-seq. This new method, MULTI-ATAC, takes advantage of the same lipid-modified oligonucleotide (LMO) system to deliver a redesigned DNA barcode oligonucleotide to the nuclear membrane. Importantly, to minimize interaction with the transposome, the barcode complex was designed to ensure no direct hybridization with Tn5 adapter sequences (Fig. S2A-B).

To first validate the efficacy and accuracy of MULTI-ATAC for pooling samples at the droplet microfluidics step, we performed a pilot experiment using peripheral blood mononuclear cells (PBMCs) from 3 unrelated donors. Nuclei from each donor were isolated separately, transposed, and uniquely barcoded, after which they were pooled and a single library was generated using the 10x Genomics scATAC-seq kit. We used deMULTIplex2 to identify doublets and assign cells to individual samples based on their MULTI-ATAC barcode counts, and then compared these classifications to those obtained by genotyping the cells using Vireo^26,27^. There was near perfect agreement between singlets identified through either method (Fig. 2A-B). The greatest degree of disagreement was in doublet classification, but we note that MULTI-ATAC-specific doublets were more similar to consensus doublets in both DoubletEnrichment scores and total fragment counts, suggesting they have a higher likelihood of being true doublets than false positives (Fig. S3A-B). We then compared these classifications against an orthogonal doublet prediction algorithm, AMULET, which is specifically designed to identify doublets in scATAC-seq data from fragment counts^28^. We note that MULTI-ATAC classifications agreed significantly with each of the other algorithms individually and in concert, and there were no Vireo-AMULET consensus doublets missed by MULTI-ATAC (Fig. S3C).

**Figure 2.**
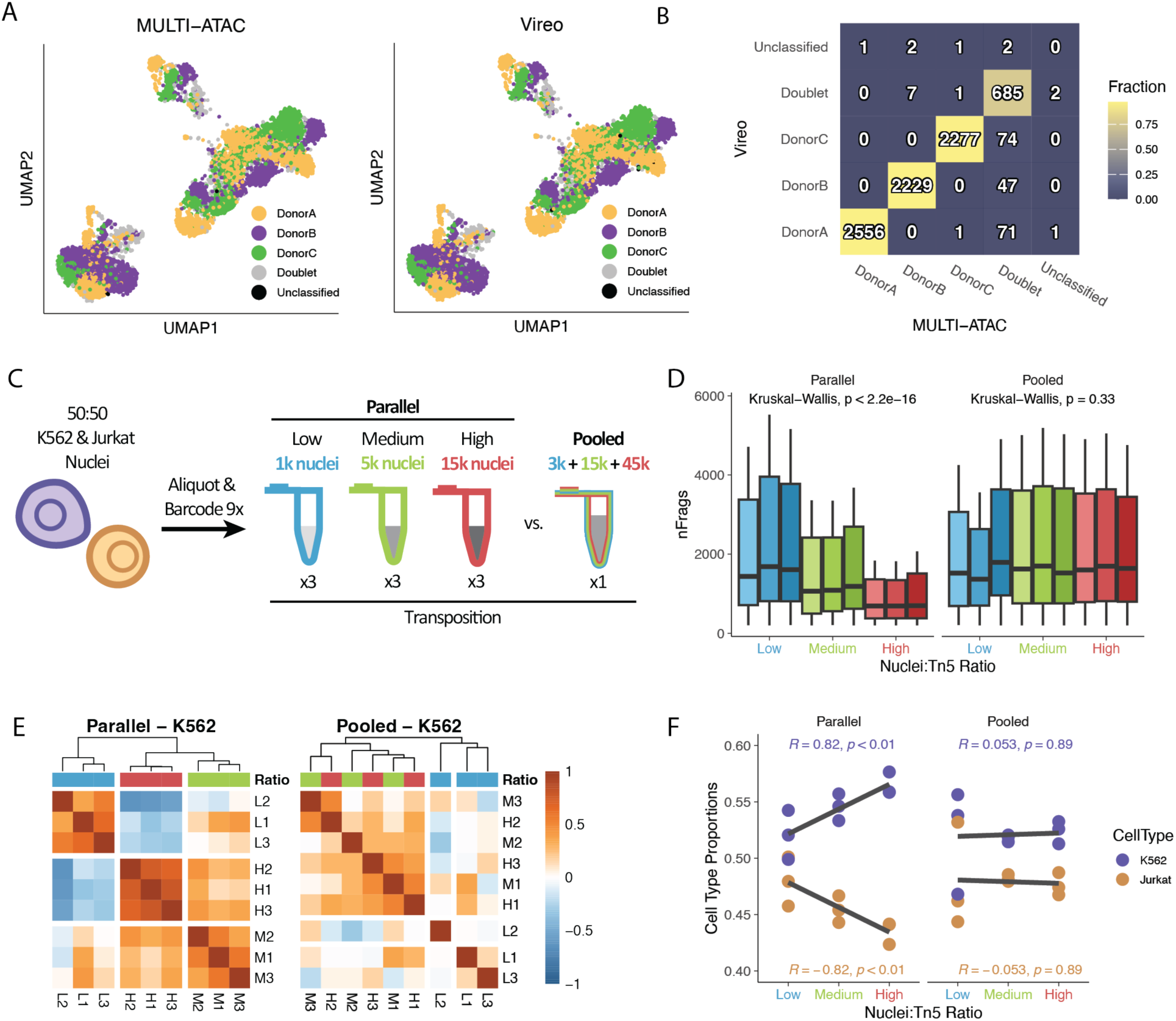
MULTI-ATAC barcoding enables pooled transposition to eliminate transposition batch size effects. A. MULTI-ATAC classifications (using deMULTIplex2) of pooled PBMC nuclei from 3 distinct donors closely matches the classifications determined by genotypic deconvolution using Vireo. B. Comparison of classification results from A) demonstrates high accuracy in singlet calling relative to genotypic deconvolution, with MULTI-ATAC/deMULTIplex2 identifying a higher rate of doublets (see Fig, S2A-C). C. Diagram of how Parallel and Pooled transposition libraries were generated from 9 uniquely-barcoded aliquots of a pool of K562 and Jurkat nuclei. D. Samples deconvolved from the Parallel library show decreasing per-nucleus fragment yield with increasing transposition batch size, whereas samples in the Pooled library all yield the same. Whisker length of boxplots shortened to 0.5 * IQR for visualization. E. Spearman correlation between per-sample means across LSI dimensions 2:30 shows strong clustering of K562 cells by transposition batch size in the Parallel library that is lost in the Pooled library. F. Similar to analysis in Fig. 1, relative proportions of K562 and Jurkat nuclei recovered per sample varied as a function of transposition batch size in the Parallel library, but were consistent across samples in the Pooled library.

### Pooled transposition with MULTI-ATAC eliminates transposition batch effects

Having validated that we can accurately assign sample identities and remove doublets using MULTI-ATAC, we next sought to investigate whether pooled transposition could ameliorate the batch effects that arise from parallel transposition reactions. To this end, we performed a “Parallel” multi-sample experiment comprising a range of nuclei yields. Specifically, we aliquoted a 50:50 mixture of K562 and Jurkat nuclei for parallel MULTI-ATAC labeling and transposition. Reactions were set up in triplicate at each of high, medium, and low nuclei:Tn5 ratios spanning the recommended range of the 10X Genomics protocol (Fig. 2C, Methods). Nuclei were then combined after transposition for library generation. In a separate library consisting of the same cell populations, we performed a “Pooled” multi-sample experiment by combining each of the 9 barcoded samples into a single pooled transposition reaction to directly assess the impact of pooled transposition on batch effects (Fig. 2C, Methods).

Mirroring our analyses of the publicly-available datasets, we observed that variable nuclei:Tn5 ratios were associated with divergent per-cell fragment yields in the Parallel library (Fig. 2D, left). In contrast, there was no density-dependent effect on fragment counts in the Pooled library (Fig. 2D, right). As demonstrated previously, variation in per-nucleus fragment counts is a covariate that influences LSI dimensionality reduction (Fig. 1E, S1B,D-E). Even when excluding the first LSI component, the 9 samples in the Parallel library clustered according to nuclei density in the reduced dimensionality space (Fig. 2E, Fig. S4A, left), a relationship that is lost when looking at cells from the Pooled library (Fig. 2E, Fig. S4A, right).

We additionally observed the expected density-dependent changes in relative proportions of each cell type in the Parallel library. Even under highly controlled conditions where equal numbers of each cell type were combined, increasing transposition batch size decreased the proportion of Jurkat nuclei from 48% to 44% and increased the proportion of K562 nuclei from 52% to 56% of the total (Fig. 2F, left). In contrast, cell type proportions remained constant across samples in the Pooled library (Fig. 2F, right). Jurkat nuclei yielded on average 36% fewer fragments than the K562 nuclei (Fig. S4B), consistent with our previous analysis that cell type proportion disparities linked to Tn5 batch size are due to the differential sensitivity of cell types to quality-control filtering (Fig. 1H).

### MULTI-ATAC empowers high sample throughput and reproducibility

Sample multiplexing approaches minimize reagent costs and improve single-cell genomics data quality through doublet detection and batch effect minimization. Beyond these benefits, multiplexing techniques provide the flexibility to execute experimental designs that are sufficiently controlled and statistically powered to derive robust conclusions. For example, high-throughput chemical screening experiments that require large numbers of individual samples (i.e., doses, replicates, and controls) are infeasible using most standard single-cell genomics workflows but become possible with the use of sample multiplexing approaches^8,29^.

To explore its utility for high-throughput single-cell genomic chemical screens, we used MULTI-ATAC to analyze the impact of perturbing the activity of 3 key epigenetic remodeling complexes (e.g., PRC2, SWI/SNF, and p300/CBP) with matched small molecule inhibitors and PROTACs in human PBMCs (Fig. 3A; Table S1). Specifically, we measured immune perturbation responses to the EZH2 inhibitor EPZ-6438 and PROTAC MS177, the SMARCA2/4 inhibitor BRM014 and PROTAC AU-15330, and the p300/CBP inhibitor GNE-781 and PROTAC dCBP-1 all in the context of T-cell activation with anti-CD3/CD28 tetrameric antibodies. Each drug was assayed at 3 doses (10nM, 100nM, and 1*µ*M) in quadruplicate along with DMSO +/- anti-CD3/CD28 antibody controls, for a total of 96 unique samples. Following 24 hours in culture, nuclei were isolated, labeled with MULTI-ATAC barcodes, and pooled for transposition prior to paired scATAC-seq and scRNA-seq profiling using the 10x Genomics Multiome platform (Fig. 3A). Notably, the same MULTI-ATAC barcoding reagents are additionally compatible with multiomic profiling^30^ (Fig S5A-B, Methods).

**Figure 3.**
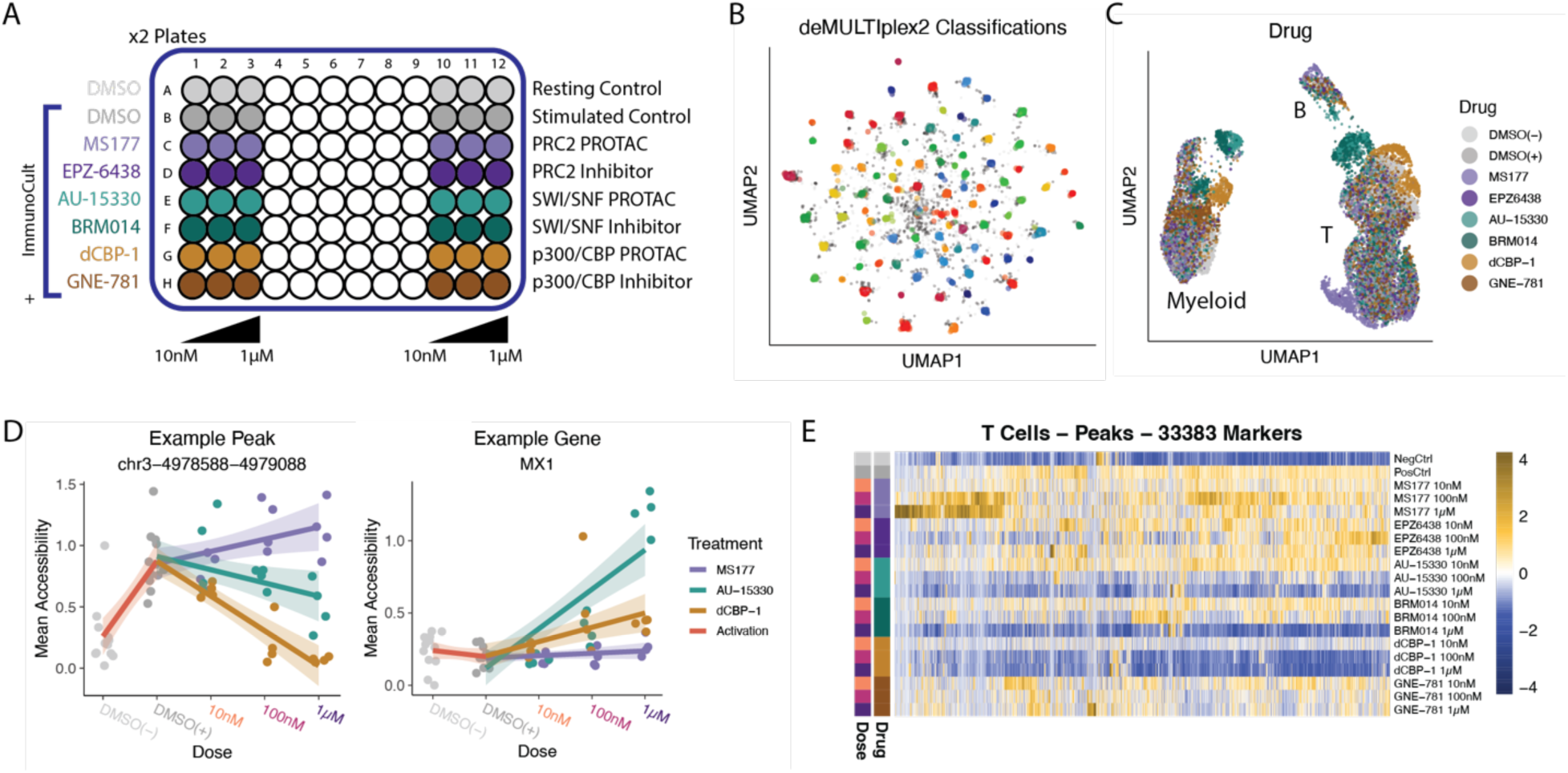
MULTI-ATAC facilitates high-throughput experimentation with reproducibility. A. Diagram of how each of two replicate 96-well plates were seeded with PBMCs and cultured with or without drugs and anti-CD3/C28 antibodies. B. UMAP embedding of MULTI-ATAC barcode counts from 1 of the 3 libraries generated, colored by which of the 96 samples each cell was classified to. C. UMAP embedding of the ATAC data generated in the Multiome experiment, colored by the drug each cell was treated with. D. Representative peak (left) and gene (right) showing how average accessibility (or expression) per cell type and replicate were used to calculate drug- or activation-responsive markers by fitting of a linear regression model. E. Heatmap of statistically significant marker peaks (p < 0.01, log2FC > 1) identified for T cells across all treatment conditions.

Following next-generation sequencing, we performed quality-control filtering and MULTI-ATAC sample demultiplexing (Fig. 3B), resulting in a final dataset of 14,233 cells. We recovered on average 148 ± 87 nuclei per tissue culture well and 609 ± 135 nuclei per drug dose, with many drugs exhibiting clear dose-dependent epigenetic reprogramming (Fig S6A-C). After unsupervised clustering and differential gene expression analysis, we identified the expected immune cell types including T cells (naïve, CD4+ and CD8+ memory, and Tregs), B cells, NK cells, and myeloid cells (monocyte and DC; Fig. S7A). Notably, a subset of treatments elicited such strong epigenetic and transcriptional responses that precluded linkage back to the subtype of origin (Fig. 3C, Fig. S6C, S7A).

The technical limitations and costs of single-cell sequencing methods typically bias study design against the inclusion of multiple biological and technical replicates. As a consequence, differential expression and accessibility analysis methods often treat individual cells as replicates or create pseudo-replicates from within individual samples, tactics which have been shown to increase the rate of false discoveries^31,32^. In contrast, using sample multiplexing to include dose regimes and true experimental replicates allows for more powerful statistical analyses that protect against artifacts (Fig. S8A-D), all increasing confidence in hypotheses emerging from experiments without increasing costs or significantly complicating workflows. We used these features of the dataset to identify high-confidence activation- and drug dose-responsive marker features for T and myeloid cells by fitting a linear regression model to the average expression or accessibility of each feature per replicate (Fig. 3D-E, Fig. S9A-F).

Effect sizes between treatments varied greatly; immune activation (particularly of T cells) almost exclusively upregulated the accessibility and expression of thousands of genes, whereas the SWI/SNF degrader AU-15330, SWI/SNF inhibitor BRM014, and p300/CBP degrader dCBP-1 mostly elicited the opposite response (Fig. 3E, Fig. S9A-F, Fig. S10A). Of note, many of the peaks that were downregulated by these drugs overlapped with the set of peaks remodeled by immune activation, predominantly reversing or inhibiting the increase in accessibility (Fig. S10B). Additionally, a large fraction of these downregulated peaks was significantly enriched for enhancer regions relative to their upregulated counterparts, particularly in myeloid cells (Fig. S10C). In contrast, the smaller subset of upregulated peaks for these drugs showed a significant enrichment for CTCF binding sites (Fig. S10D). Myeloid cells were particularly sensitive to this effect, perhaps in part because a greater fraction of the accessible chromatin in these cells was associated with annotated distal enhancer regions (Fig. S10E-F). Because CTCF acts to insulate regions of the genome as topologically-associated domains to promote enhancer-gene interactions, the concurrent loss of enhancer accessibility and increase in CTCF site accessibility may reflect a mechanism by which these drugs impact 3D chromatin organization.

### Epigenetic perturbations elicit drug- and cell-type specific effects

We next analyzed the differential impact of drugs targeting the same complex by direct inhibition or degradation. To visualize the overlapping and varied impacts of these drugs on immune cells we developed a two-dimensional scoring system that decomposed the drug effects into two components reflecting influences on immune activation versus all other effects on chromatin accessibility (Fig. 4A, Methods). We then used this scoring system to compare PROTAC-inhibitor pairs across a 3-order of magnitude dose regime (Fig. 4A). The analysis revealed divergent responses in distinct immune cell populations linked to both drug target and mechanism of action. For example, we found that SWI/SNF disruption was highly dose-responsive and that equimolar treatments with either the PROTAC AU-15330 or inhibitor BRM014 elicited similar responses in T and myeloid cells (Fig 4A, center). By contrast, the PROTAC dCBP-1 produced a much stronger response in both T and myeloid cells than the inhibitor GNE-781 from which it is derived, supporting previous findings about the potency of p300/CBP degradation over inhibition^33^ (Fig. 4A, right). Finally, we observed a cell-type-specific ‘bell-shaped’ dose-response pattern in T cells treated with the EZH2 PROTAC, MS177, where the 100nM dose induced increased activation before dropping back down at 1*µ*M (Fig. 4A, left). This result was not observed in cells treated with the EZH2 inhibitor EPZ-6438, which exhibited little overall phenotype. Notably, this trend coincides with a set of “amplified” activation-associated peaks noted for this drug in T cells, lending credence to this scoring metric (Fig. 3E, Fig. S10B).

**Figure 4.**
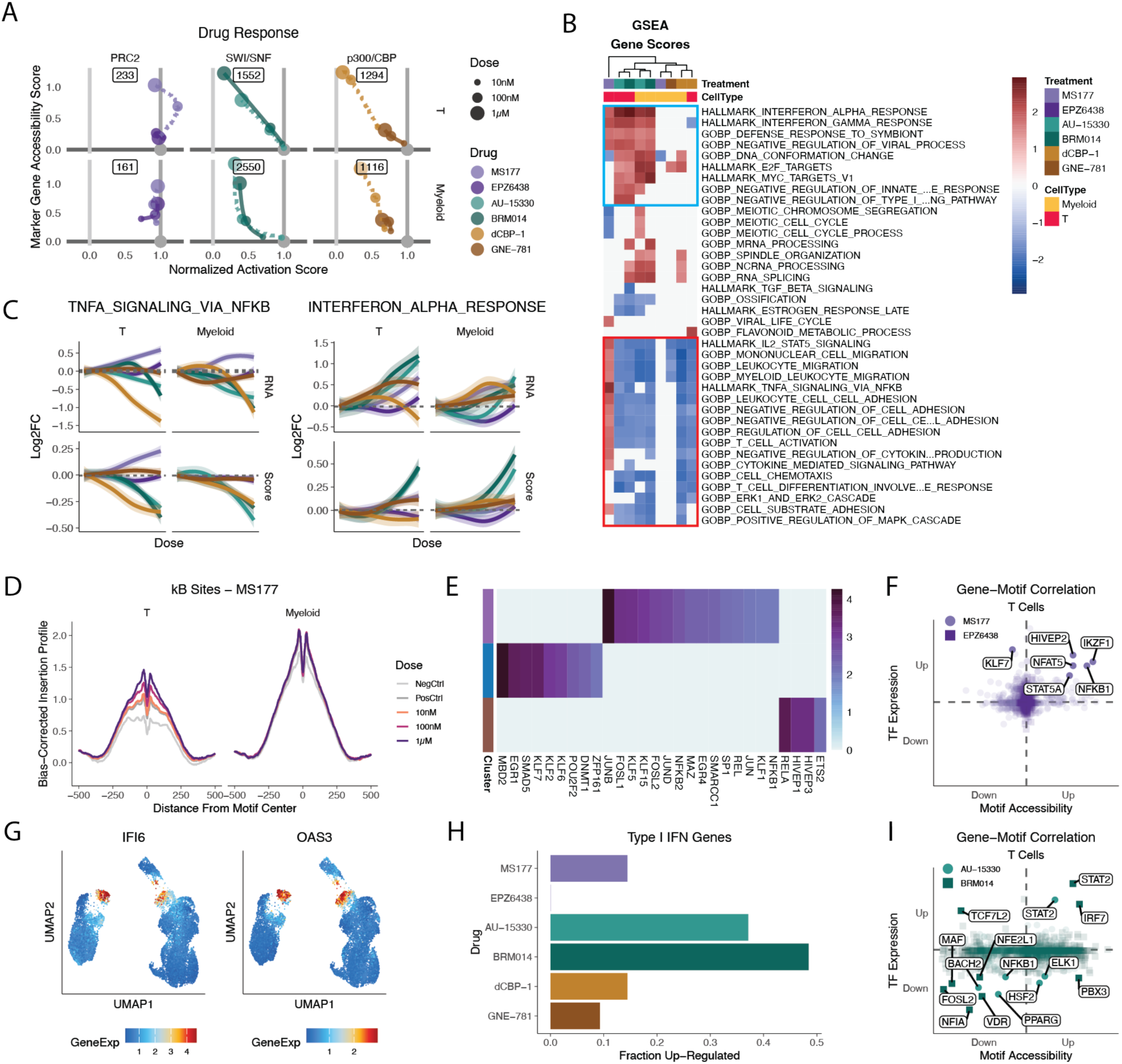
Drug- and cell type-specific effects of epigenetic perturbation. A. Two-component drug response analysis; the X-axis scores each drug dose by its relative activation compared to controls using activation-associated marker gene scores, while the Y-axis scores each drug dose on the accessibility of drug-responsive marker genes not associated with activation. Solid lines show the dose-response trajectory of inhibitors, whereas dashed lines show the trajectories of PROTACs. Inset values show the number of drug-responsive marker genes used to generate the Y-axis scores. See Methods for more details. B. Gene set enrichment analysis (GSEA) for each drug and cell type of markers gene accessibility scores ordered by statistical significance and negative vs positive slope. Statistically significant terms (p.adj. < 0.01) are colored by normalized enrichment score (NES). Red box – gene sets involved in immune cell activation; blue box – gene sets involved in type I interferon response. C. Gene set expression across increasing drug dose; log2 fold-changes in expression or accessibility of each gene at each drug dose were calculated relative to the activated controls, and then plotted as a function of dose. Trendlines plotted per drug via LOESS smoothing with span = 1.5. D. NF-κB motif footprinting in control and MS177-treated T and myeloid cells. E. Significantly enriched TF motifs (p.adj. < 0.01) across 3 clusters of MS177-responsive peaks in T cells (see Fig. S11C). Heatmap colored by -log10(p.adj.). F. Correlation of TF motif accessibility and TF RNA expression. Axes represent increasing statistical significance of negative/positive relationship with MS177 or EPZ-6438 dose. Solid, annotated points are statistically significant (p < 0.01) in both modalities. G. UMAP embeddings showing imputed RNA expression values for two representative Type I Interferon response genes. H. Fraction of Halllmark Interferon Alpha Response gene set upregulated in accessibility and/or expression across any cell type. I. Same as F) but for AU-15330 and BRM014.

To further contextualize these results, we investigated drug-specific effects on immune cells using pathway analysis. We ranked genes by the strength and direction of their response to drug treatment (both in terms of accessibility and RNA expression) and performed gene set enrichment analysis^34^ on the ranked lists (Fig. 4B, Fig. S11A). As expected, terms related to immune activation and differentiation were downregulated specifically in the dCBP-1, AU-15330, and BRM014 samples that also exhibited the greatest inhibition of immune activation. Notably, many of these same terms were upregulated in MS177-treated T cells (Fig. 4B & Fig. S11A, red box), underscoring that this drug may uniquely amplify the activation state of the cells.

Of the gene sets upregulated by MS177 in T cells, the most significantly enriched is TNFα signaling via NF-κB. In aggregate, these genes exhibited a dose-dependent increase in RNA expression relative to positive controls in both T cells and myeloid cells, whereas their gene accessibility only increased noticeably in T cells (Fig. 4C). We hypothesized that this deviation between RNA and ATAC data was due to myeloid cells having higher baseline expression and gene accessibility of these genes relative to T cells (Fig. S11B, left). To test this notion, we profiled the accessibility of NF-κB binding sites genome-wide and observed that while MS177 treatment increased the accessibility of these sites in T cells, in myeloid cells these sites were highly accessible at baseline and insensitive to treatment despite the increase in target gene expression (Fig. 4D).

Beyond cell-type-specific chromatin remodeling near NF-κB binding sites, hierarchical clustering of MS177 and activation marker peaks in T cells revealed that most MS177-responsive peaks seemed to cluster into three main groups (Fig. S11C, brown, purple, blue): two that increased in accessibility sharply with MS177 dose and were unrelated to activation, while the third included activation-associated peaks and reached maximum accessibility at the 100 nM dose and dropped thereafter, mirroring the activation score analysis. These peak sets were strongly enriched with binding sites for NF-κB family members, AP-1 family members, and other transcription factors critical to T cell function (Fig. 4E)^35^. To better ascertain which exact transcription factors may drive the response to MS177, we looked specifically at factors whose RNA expression and motif accessibility both increased in response to MS177 treatment. This analysis highlighted a variety of genes involved in T-cell activation, differentiation, and exhaustion such as NFKB1, NFAT5, STAT5A, HIVEP2, and IKZF1 (Fig. 4F)^36–39^.

We next sought to characterize the epigenomic and transcriptomics responses to SWI/SNF and p300/CBP inhibition in human PBMCs. While SWI/SNF- and p300/CBP-targeting drugs largely decreased both chromatin accessibility and gene expression relative to activated controls (Fig. S9A-F, Fig. S10A), these samples exhibited enrichment for gene sets associated with type I interferon signaling, the innate immunity pathway largely responsible for mounting early responses to pathogenic infection (Fig. 4B & Fig. S11A, blue box)^40–42^. In particular, the SWI/SNF-targeting drugs AU-15330 and BRM014 demonstrated a clear and dose-dependent increase in both the expression and accessibility of interferon-stimulated genes (ISGs) and upstream regulators, irrespective of cell type (Fig. 4C,G-H, Fig. S11B, Fig. S12A). Specifically, we observed upregulation of terms and genes pertaining to antiviral response and detection of foreign RNA and DNA (Fig. S12A-B). In line with these results, we observed that these drugs induce concurrent increases in expression and motif accessibility for transcription factors involved in interferon signaling, notably IRF7 and STAT2 (Fig 4I). Finally, other upregulated terms related to transcription, splicing, and DNA-nucleosome interactions, all of which exhibited increased accessibility without a corresponding increase in RNA expression (Fig. 4B, Fig. S11B, Fig. S12B). Among these dysregulated genes were the replication-dependent histones — for instance, the HIST1 gene cluster on chr6 showed a dose-dependent increase in accessibility that was most pronounced in the SWI/SNF-targeting drugs (Fig. S13A-B). While the cause of this is unknown, one possible explanation is that SWI/SNF inhibition in particular prevents expression of genes necessary for progression through the cell cycle^43,44^.

## Discussion

Despite efforts to increase the scalability of scATAC-seq methods using multiplexing or combinatorial indexing, enzymatic transposition remains a limiting step, requiring that many separate parallel reactions be run simultaneously. Perhaps more concerningly, we identified previously unappreciated technical batch variation in publicly available datasets that use parallel transposition reactions that can be traced back to variable nuclei inputs across reactions – a finding we confirm experimentally. While this type of batch effect is not wholly unexpected considering similar findings in bulk ATAC-seq data, it is either rarely addressed or thought to be removed during pre-processing steps of typical analysis pipelines. Instead, we demonstrate that transposition batch effects are readily detectable across many publicly available datasets, are not easily removed using current data processing best practices, and impact downstream biological interpretation.

A key finding is that transposition batch size biases compositional analyses for or against certain cell types. Variation in cell type composition between individuals or in response to treatments can be biologically impactful and is thus important to understand and report accurately. For example, a decrease in cancer cells and increase in infiltrating immune cells in response to a new immunotherapy drug would be an indicator of clinical response. We find that variation in nuclei per sample can generate precisely this type of shifts in data. When aggregated and averaged across dozens of transposition batches such as in some sci-ATAC-seq3 datasets, these effects may become less severe. However, when the number of transposition reactions per sample is low or a sample is transposed in a single reaction, common for droplet microfluidics workflows, the risk of analyses being influenced by nuclei counts and per-nucleus fragment yield is significant.

To overcome this technical hurdle, we developed MULTI-ATAC, a method for labeling nuclei with sample-specific DNA barcodes that can be sequenced alongside scATAC-seq libraries. Using genotypically-distinct donor samples, we demonstrate the ability of MULTI-ATAC barcoding to reliably and accurately assign sample identities to nuclei pooled during library preparation. While almost no cells were misassigned to the wrong sample-of-origin, we did note increased rates of doublet-calling compared to two *in silico* methods. While we cannot rule out if these were false-positive doublet assignments, we observed that these particular cells shared similarities with bona fide doublets. Additionally, whereas the two other classification methods, AMULET & Vireo, rely on the sequenced chromatin fragments as input to classify each cell, MULTI-ATAC barcode counts represent an orthogonal modality that does not necessarily depend on per-nucleus ATAC data quality. It is therefore possible that MULTI-ATAC classifications are closest to ground truth.

We next utilized MULTI-ATAC barcoding to explicitly demonstrate how pooled transposition removes batch effects. We processed 9 samples, either in parallel or in a pooled format, at different nucleus-to-Tn5 ratios spanning the range recommended by commercially available scATAC-seq kits from 10X Genomics. By quantifying batch effects at the levels of data quality, clustering, and sample composition, we found that pooled processing enabled by MULTI-ATAC eliminates batch effects present in the parallel-processed samples. These findings demonstrate that realistic variability in transposition conditions could easily impact sample comparisons within and between individual experiments if inputs are not carefully controlled.

Finally, to demonstrate the scope of experimental designs made possible by MULTI-ATAC, we performed a 96-plex drug screen of epigenetic inhibitors and degraders in human immune cells. Single-cell drug assays are typically challenging and expensive to perform due to the inherently high number of samples, and researchers must often compromise either the number of replicates or the number of doses assayed. The facility of MULTI-ATAC barcoding and pooled transposition means the number of samples one can assay is limited primarily by the nuclei isolation step and the number of unique MULTI-ATAC barcode sequences one has. With MULTI-ATAC we were able to include both a 3 order-of-magnitude dose regime as well as four replicates for each dose of 6 different drugs. This enabled downstream analyses that are robust to technical and biological variation between replicates without inflating p-values from treating each cell as an individual replicate.

Analysis of the drug responses revealed numerous drug-, target-, and cell type-specific effects. Most apparent was the differential response to the EZH2 degrader MS177 and inhibitor EPZ-6438. Specifically, we found that the EZH2 inhibitor EPZ-6438 showed little impact on the transcriptomes and epigenomes of the cells in culture at any dose. This is likely because the primary mechanism of clearance of H3K27me3, the repressive histone modification catalyzed by EZH2/PRC2, has been shown to be replicative dilution^45^. We would therefore expect that a longer culture period and multiple population doublings would be required for EPZ-6438 to start exhibiting effects.

By contrast, the EZH2 degrader MS177 very potently altered the T and myeloid cells, inducing increased expression and/or accessibility of NF-κB associated genes and motifs. NF-κB signaling is a known contributor to signaling downstream of TCR activation, which partially explains the augmented T cell activation exhibited by the 100 nM dose of MS177. The mechanistic relationship between MS177 treatment and NF-κB signaling is not yet understood; however, several avenues for further investigation are evident from the data. For instance, a pair of studies have demonstrated direct physical interactions between EZH2 and NF-κB factors that contribute to transcriptional regulation independently of methyltransferase activity^46,47^. NF-κB pathways invoke degradation of downstream mediators as part of the signaling cascade; therefore, one hypothesis is that MS177 amplifies NF-κB signaling activity by concomitantly degrading a negative NF-κB regulator associated with EZH2. Another notable finding regarding MS177 treatment is the upregulation of the IKZF1/Ikaros and IKZF3/Aiolos transcription factors, which are important regulators of lymphocyte function and development. Intriguingly, these proteins have been identified as neo-substrates of the CRBN ubiquitin ligase that is recruited by MS177^33,48–51^, and Ikaros has been shown to both associate with PRC2 and mediate T cell exhaustion through repression of AP-1, NFAT, and NF-κB target genes^39,52^. Taken together, it is possible that MS177 exerts these effects through off-target degradation of IKZF1/IKZF3, leading to upregulation of downstream targets related to T cell activation.

The drugs targeting the SWI/SNF nucleosome remodeling complex and p300/CBP histone acetyltransferases primarily seemed to inhibit lymphocyte activation and led to variable decreases in both chromatin accessibility and gene expression. Despite this, two groups of gene sets exhibited pronounced upregulation during pathway analysis. Genes related to cell cycle and RNA processing became more accessible but were not upregulated transcriptionally; simultaneously, a pronounced type I interferon response was induced. Multiple studies have demonstrated that epigenetic dysregulation can stimulate a type I interferon response through the de-repression of human endogenous retrovirus (ERV) and other retrotransposons, and that this is likely to contribute to age-related inflammation and disease^53–56,56–61^. More recently, mutations, deficiencies, and perturbations of several different SWI/SNF-family proteins have been shown to induce cell-intrinsic type I interferon responses in cancer cells that can improve the response of tumors to immune checkpoint blockade^56,58,62–64^. In these studies, interferon signaling is traced back to numerous mechanisms including ERV expression, R-loop formation, and excess cytoplasmic ssDNA production, with both DNA- and RNA-sensing pathways implicated. Depletion of H1 linker histones has also been shown to induce interferon signaling, providing a possible link to issues with cell cycle progression^65–67^. The breadth of evidence supporting a more general mechanism linking innate immune activation to perturbed chromatin organization indicates this to be an exciting area for future investigation.

While MULTI-ATAC barcoding stands to greatly improve scATAC-seq workflows by allowing pooled transposition, we note that other workflow bottlenecks still impede large scale experiments. Barcoding itself is fast and can be done at various scales without significant optimization. Nuclei isolation, however, is a step that all investigators must contend with and optimize for their sample type. Scaling up to many samples carries inherent risk of introducing batch effect if lysis times are not properly controlled. However, we note that the ability to include many replicates enables hedging against such challenges.

Finally, Tn5 transposition has been harnessed in a growing variety of sequencing assays, including mitochondrial DNA sequencing, proteomics, profiling of DNA-binding proteins, and 3D chromatin mapping^68–73^. Because most depend on capturing transposed fragments on the 10x Genomics platform, we hypothesize that, perhaps with only minor protocol adjustments, MULTI-ATAC barcoding could be successfully extended to many of these methods as well to great effect.

## Methods

### Design of MULTI-ATAC protocol and oligonucleotides

LMO-based barcoding of nuclei was adapted from MULTI-seq^1^ using stand LMO Anchor and Co-Anchor components available from MilliporeSigma. To mimic gDNA fragments and enable single-cell barcoding by 10x Genomics scATAC-seq kits or similar technologies, the 5’ end of the ssDNA barcode begins with the full Nextera R1 sequence. This is followed by a unique molecular identifier (UMI) of 8 random bases (N’s), a predetermined 8-base sample-specific barcode (X’s), and a TruSeq R2 sequence to enable barcodes to be separately amplified from ATAC fragments. At the 3’ end is the TruSeq Small RNA R2 sequence which hybridizes to the LMO Anchor. The inclusion of the internal TruSeq R2 site for library amplification was intended to protect against degradation of the primer site by possible 3’-5’ exonuclease activity during in-GEM linear PCR, but this was not explicitly tested.

The 5’-3’ orientation of the ssDNA barcode prevents direct hybridization to the Nextera adapter oligos in the Tn5 transposome, and is not immediately compatible with the orientation of the capture oligos employed by 10x Genomics in v1 and v2 scATAC-seq kits. To overcome this, a Barcode Extension primer is pre-annealed to the MULTI-ATAC barcode before labeling. This primer is extended during the initial gap-fill reaction in droplets which produces the complement strand needed for in-GEM capture and linear amplification of barcode oligos alongside ATAC fragments.

Because MULTI-ATAC barcodes are similar in size to the smallest ATAC fragments, they cannot be size-separated during scATAC-seq library preparation without loss of ATAC fragments. Thus, the barcode library is generated from a 1*µ*L aliquot that is taken from each scATAC-seq library prior to the Sample Index PCR step. This aliquot is amplified in a separate sample indexing PCR reaction using the same SI-PCR-B Fwd primer (ordered separately to control concentration) as the scATAC-seq libraries and a custom TruSeq Rev primer with a unique library-specific i7 index.

MULTI-ATAC barcode: 5’-TCGTCGGCAGCGTCAGATGTGTATAAGAGACAG**NNNNNNNNXXXXXXXX**AGATCG GAAGAGCACACGTCTGAACTCCAGTCACCCTTGGCACCCGAGAATTCCA-3’

Barcode Extension primer: 5’-GTGACTGGAGTTCAGACGTGTGC-3’

TruSeq-# primer: 5’-CAAGCAGAAGACGGCATACGAGAT**XXXXXX**GTGACTGGAGTTCAGACGTGTGCTCTTCCGATCT-3’

SI-PCR-B primer: 5’-AATGATACGGCGACCACCGAGA-3’

### Cell culture

Cryopreserved PBMCs were thawed in a 37°C water bath before gently transferring to a 50mL conical vial and adding 10x volume (10-20mL) of RPMI 1640 culture media. Cells were pelleted at 400rcf, 4°C, for 4 minutes, before resuspending in RPMI 1640 media supplemented with 10% fetal bovine serum and 1% penicillin-streptomycin and seeding in an ultra-low attachment 10cm culture dish. PBMCs were allowed to incubate at rest for 24 hours prior to subsequent experimental steps. K562 and Jurkat cells were thawed in a 37°C water bath, plated at 1M/mL, and cultured for several passages in RPMI 1640 media, supplemented with GlutaMAX, 10% fetal bovine serum, and 1% penicillin-streptomycin. All cells were incubated at 37°C, 5% CO2.

### Nuclei isolation

Unless noted otherwise, cell suspensions were first washed once with chilled PBS. 500k cells per sample were aliquoted into 1.5mL Eppendorf tubes and pelleted at 300rcf, 4°C, for 4 minutes. Cells were resuspended in 100 *µ*L of chilled Lysis Buffer (10 mM Tris-HCl pH 7.4, 10mM NaCl, 3mM MgCl2, 0.1% Tween-20, 0.1% Nonidet P40 Substitute, 0.01% Digitonin, 2% BSA in nuclease-free water), mixed, and incubated 5 minutes on ice. Then, 1 mL Wash Buffer (10 mM Tris-HCl pH 7.4, 10mM NaCl, 3mM MgCl2, 0.1% Tween-20, 2% BSA in nuclease-free water) was added and mixed. Nuclei were pelleted at 500rcf, 4°C, for 4 minutes and then resuspended in chilled PBS.

### MULTI-ATAC barcoding

Unless noted otherwise, MULTI-ATAC barcode complexes were assembled by combining LMO Anchor, barcodes, and BE primer in a 2:1:2 molar ratio in nuclease-free water. We found that including excess LMO Anchor and BE Primer improved barcode capture (data not shown). Isolated nuclei were adjusted to a concentration of 750-1000 nuclei per *µ*L. Assembled barcode complex was added to each nuclei suspension at 10nM, 25nM, or 50nM labeling concentration, followed by mixing by vortex pulse or pipette and incubation on ice. After 5 minutes, LMO Co-Anchor was added at twice the concentration of the full barcode complex (to account for excess LMO Anchor), mixed, and incubated another 5 minutes on ice. Barcoding was quenched by addition of 1.2mL 2% BSA in PBS. Barcoded nuclei were pelleted at 500rcf, 4°C, for 4 minutes, then resuspended in 100-200*µ*L 2% BSA in PBS for counting and pooling with other samples.

### Multi-donor pilot experiment

Three distinct vials of PBMCs from different donors and vendors were thawed and cultured as described previously. After 24 hours, each batch of PBMCs was divided into multiple 500k cell aliquots for nuclei isolation as described previously. Isolated nuclei from each donor were concentrated to 7.5k nuclei/*µ*L, from which 4 *µ*L were added to PCR strip tubes containing 26 *µ*L of transposition mix (15 *µ*L 2X Tagment DNA Buffer, 5.9 *µ*L PBS, 0.3 *µ*L 10% Tween-20, 0.3 *µ*L 1% Digitonin, 1.5 *µ*L Tagment DNA Enzyme 1, 3 *µ*L nuclease free water). The tubes were incubated at 37°C in a thermocycler for 1 hour. Transposed nuclei were barcoded as described before except that barcode complexes were assembled at 1:1:1 molar ratio. Both barcode complex and LMO Co-Anchor were added at a final concentration of 25 nM. Barcoded, transposed nuclei from each donor were then pooled and resuspended to a density of 1k/*µ*L in ATAC Buffer B before proceeding with scATAC-seq library generation with the 10x Genomics Single Cell ATAC v1.1 kit.

### Parallel vs pooled transposition batch effect experiment

Nuclei were isolated from K562 and Jurkat cells, counted, and pooled at equal numbers. 9 aliquots were drawn from this pool for MULTI-ATAC barcoding as described previously. These 9 aliquots were diluted to 200 nuclei/*µ*L, 1k nuclei/*µ*L, or 3k nuclei/*µ*L, and then 9 parallel transpositions were set up, combining 10 *µ*L of each nuclei mixture with 20 *µ*L transposition mix (15 *µ*L 2X Tagment DNA Buffer, 0.3 *µ*L 10% Tween-20, 0.3 *µ*L 1% Digitonin, 1.5 *µ*L Tagment DNA Enzyme 1, 2.9 *µ*L nuclease free water). Simultaneously, the same ratios of each of the 9 barcoded aliquots were combined and 45 *µ*L of this mixture was added to 90 *µ*L of transposition mix. The 9 parallel transposition tubes and 1 pooled transposition tube were all incubated at 37°C in a thermocycler for 1 hour, after which the parallel tubes were pooled. Both barcoded, transposed nuclei suspensions were then counted and resuspended to a density of 1k nuclei/*µ*L in a 1:2 mixture of 1X Nuclei Buffer and ATAC Buffer B before proceeding with scATAC-seq library generation with the 10x Genomics Single Cell ATAC v2 kit.

### Multiome pilot experiment

Mouse hepatocytes were isolated by a two-step perfusion technique. Briefly, mouse was anesthetized by isoflurane (Piramal Critical Care). Mouse liver and heart were exposed by opening the abdomen and cutting the diaphragm away. The portal vein was cut and immediately the *inferior vena cava* was cannulated via the right atrium with a 22-gauge catheter (Exel International, 26746). Liver was perfused with liver perfusion medium (Gibco, 17701038) for 3’ and then with liver digest medium (Gibco, 17703034) for 7’ using a peristaltic pump (Gilson, Minipuls 3). Pump was set to 4.4 mL/min and solutions were kept at 37°C. After perfusion the liver was dissected out, placed in a petri dish with hepatocyte plating medium (DME H21 [high glucose, UCSF Cell Culture Facility, CCFAA005-066R02] supplemented with 1x PenStrep solution [UCSF Cell Culture Facility, CCFGK004-066M02], 1x Insulin-Transferrin-Selenium solution [GIBCO, 41400-045] and 5% Fetal Bovine Serum [UCSF Cell Culture Facility, CCFAP002-061J02]) and cut into small pieces. Liver fragments were passed through a sterile piece of gauze. Hepatocytes were separated from non-parenchymal cells by centrifugation through 50% isotonic Percoll (Cytiva, 17-0891-01) solution in HAMS/DMEM (1 packet Hams F12 [GIBCO, 21700-075], 1 packet DMEM [GIBCO, 12800-017], 4.875 g sodium bicarbonate, 20 mL of a 1M HEPES pH 7.4, 20 mL of a 100X Pen/Strep solution, 2 L H_2_O) at 169 g for 15’. Isolated hepatocytes were frozen in BAMBANKER (GC LYMPHOTEC, CS-02-001) and stored at -80°C.

On the day of the experiment, frozen hepatocytes were thawed, washed with PBS (Gibco, 10010-023) and fixed in 1% PFA (Electron Microscopy Sciences, 15714-S) for 10 min at RT. Fixation was quenched by addition of glycine (125 mM final concentration) and washed with cold PBS supplemented with 1% BSA (Sigma, A1953). Hepatocytes were next permeabilized by resuspending 0.5 million fixed cells in 100 μL of lysis solution (0.5% n-Dodecyl β-D-maltoside, 45 mM NaCl, 10 mM Tris-HCl pH 8.0, 5 mM MgCl_2_, 10% dimethylformamide, 1U/*µ*L Protector RNase inhibitor [MilliporeSigma, 3335399001]) and incubated on ice for 5 minutes. Permeabilization was stopped by adding 1 mL of wash buffer (45 mM NaCl, 10 mM Tris-HCl pH 8.0, 5 mM MgCl_2_, 1% BSA, 1U/*µ*L Protector RNase inhibitor [MilliporeSigma, 3335399001]). Next, fixed, permeabilized cells were barcoded with both MULTI-seq and MULTI-ATAC reagents. LMO Anchor was assembled into complex with MULTI-seq barcodes (2:1 ratio) or with MULTI-ATAC barcodes and BE primer (2:1:2 ratio). Cells were divided into 5 aliquots, two were labeled with MULTI-ATAC barcodes as described, two were labeled with MULTI-seq barcodes following the same protocol, and the fifth aliquot was left unlabeled as a control. All 5 aliquots were pooled, resuspended in 1X Nuclei Buffer and adjusted to 5k cells/*µ*L for processing with the 10x Genomics Single Cell Multiome ATAC + Gene Expression v1 kit.

### Multiome epigenomic drug screen

PBMCs from a single donor were thawed and cultured as described. After resting for 24 hours, non-adherent cells and media were transferred to a 50 mL conical vial. Pre-warmed TrypLE was added to culture dish and incubated 2 minutes at 37°C to lift remaining cells before also transferring to conical vial. Cells were pelleted at 400rcf, RT, for 4 minutes, and resuspended in PBS to count and assess viability. After, cells were resuspended in media (RPMI 1640, 10% FBS, 1% Pen/Strep) to 1k cells/*µ*L. 192.5 *µ*L of cell suspension were deposited into each well of the outermost 6 columns of two 96-well ultra-low attachment round-bottom plates. To each well was then added 2.5 *µ*L of 80X drug-media solution or 2.5 *µ*L of DMSO-media solution, and 5 *µ*L of ImmunoCult anti-CD3/CD28 antibodies or equivalent volume of PBS. All wells were gently pipette-mixed 5X with a multichannel p200 set to 150 *µ*L. Plates were returned to the incubator and cultured 24 hours.

The following day, cells were gently pipette mixed to resuspend and then pelleted at 400rcf, 4°C, for 5 minutes. Media was carefully aspirated and pellets were resuspended in 100 *µ*L 2% BSA in PBS, before transferring cells to a set of new 96-well ultra-low attachment round-bottom plates on ice. To recover remaining adhered cells, 100 *µ*L of pre-warmed TrypLE was added, followed by 2 minute incubation at 37°C, and transfer of the full 100 *µ*L to the new plates on ice. 100 *µ*L from each well was aliquoted into a new set of standard 96-well round-bottom plates and pelted at 400rcf, 4°C, for 5 minutes. 95 *µ*L were carefully removed from each well. Then pellets were resuspended in 45 *µ*L chilled lysis buffer (10 mM Tris-HCl pH 7.4, 10mM NaCl, 3mM MgCl2, 0.1% Tween-20, 0.1% Nonidet P40 Substitute, 0.01% Digitonin, 1 mM DTT, 1 U/*µ*L Protector RNase inhibitor (MilliporeSigma, 3335399001), 1% BSA in nuclease-free water) and pipette-mixed 3X. Lysis was allowed to proceed 2.5 minutes, with the timer being initiated after addition of buffer to the first column. At the end of incubation, 150 *µ*L wash buffer (10 mM Tris-HCl pH 7.4, 10mM NaCl, 3mM MgCl2, 0.1% Tween-20, 1 mM DTT, 1 U/*µ*L Protector RNase inhibitor (MilliporeSigma, 3335399001), 1% BSA in nuclease-free water) was added without mixing. Plates were pelleted at 600rcf, 4°C, for 5 minutes, after which 195 *µ*L of supernatant was carefully removed and discarded.

Pellets were resuspended in 95 *µ*L chilled PBS, after which 50 *µ*L of one of each 96 unique pre-assembled 75 nM MULTI-ATAC barcode complexes (2:1:2 molar ratio) was added to each well and gently pipette-mixed, for a final labeling concentration of 25 nM. Plates were left on ice for 5 minutes, before addition of 50 *µ*L of 200nM LMO Co-Anchor, gentle pipette-mixing, and another 5 minutes on ice. Plates were pelleted at 600rcf, 4°C, for 5 minutes, before aspirating 195 *µ*L of supernatant and resuspending each well in 195 *µ*L chilled 2% BSA in PBS to quench labeling.

100 *µ*L from each well were pooled by row, pelleted, and resuspended in 50 *µ*L 1X Nuclei Buffer for counting. The row pools were merged together, adjusted to 3-5k nuclei/*µ*L, and processed with the 10x Genomics Single Cell Multiome ATAC + Gene Expression v1 kit.

During analysis, we noted a significant separation in the UMAP embedding between cells originating from the left and right side of the 96-well plates they were cultured and lysed in. Deeper inspection of the data revealed that LSI component 4 seemed to capture the bulk of this variance. Additionally, marker analysis between matched “left-side” and “right-side” cells predominantly showed differences in promoter accessibility (data not shown), which correlates with slight but statistically significant differences in QC metrics. Therefore, this variance was deemed to likely be a technical artifact from either culture or lysis, and this component was excluded from downstream embedding. Importantly, this decision primarily affected visualization and did not influence later marker analyses.

### scATAC-seq library preparation

Unless otherwise noted, pooled, barcoded nuclei were transposed and subsequently processed into scATAC-seq libraries according to manufacturer’s recommendations (10x Genomics), with only minor modifications. Briefly, at step 3.2o, a 1 *µ*L aliquot is taken from each individual library to be used in producing accompanying MULTI-ATAC barcode libraries. This left only 39 *µ*L to be carried into the subsequent Sample Index PCR reactions (step 4.1), where we also exchanged the SI-PCR Primer B with an equivalent volume of a 100 *µ*M SI-PCR-B primer with the same sequence, ordered separately (IDT).

### Multiome library preparation

Barcoded nuclei or fixed permeabilized cells were transposed and subsequently processed into paired single-cell GEX and ATAC libraries according to manufacturer’s recommendations (10x Genomics), with only minor modification. Briefly, after Pre-Amplification PCR (step 4.2) completed, a 1 *µ*L aliquot was taken from each PCR reaction to be used in producing accompanying MULTI-ATAC barcode libraries.

### MULTI-ATAC barcode library preparation

1 *µ*L aliquots from each scATAC-seq or Multiome library preparation were taken at the appropriate step (see above) and incorporated into a PCR reaction with 2.5 *µ*L 10*µ*M SI-PCR-B primer, 2.5 *µ*L TruSeq-# indexing primer, 26.25 *µ*L Kapa HiFi HotStart ReadyMix, and 17.75 *µ*L nuclease-free water. The reaction was run with the following protocol: 1. 95°C/5:00, 2. 98°C/0:20, 3. 67°C/0:30, 4. 72°C/0:20, 5. repeat steps 2-4 x13, 6. 72°C/1:00, 7. 4°C/hold. Afterwards, 100 *µ*L SPRIselect were added, pipette-mixed 10x, and incubated 5’ at RT. Tubes were placed on a magnet rack and beads washed with two successive additions of 200 *µ*L fresh 80% EtOH, with 30” pauses between. EtOH was aspirated and libraries were eluted from beads for 2’ at RT in 20 *µ*L Buffer EB.

### MULTI-seq barcode library preparation

MULTI-seq barcodes were prepared for the Multiome Pilot Experiment similarly to as described previously^1^, with minor modifications. 10 *µ*L of Pre-Amplification SPRI Cleanup product (step 4.3p of Multiome protocol) were transferred into a fresh PCR strip tube, to which 40 *µ*L Buffer EB were added. 30 *µ*L SPRIselect reagent (0.6X) were added, pipette mixed, and incubated 5’ at RT. Strip tube was placed on a magnet rack, and the supernatant containing MULTI-seq barcodes was transferred to a fresh 1.5 mL tube. 130 *µ*L SPRIselect (3.2X) and 90 *µ*L fresh isopropanol (1.8X) were added to this supernatant, mixed, and incubated 5’ at RT. After placing on magnet rack and discarding supernatant, MULTI-seq library preparation was carried on from step 15 as normal.

### Sequencing & library pre-processing

All scATAC-seq and Multiome libraries were sequenced on NovaSeq 6000 SP, NovaSeq 6000 S4, or NovaSeq X 10B flow cell lanes according to manufacturer’s recommendations (10x Genomics). Briefly, for scATAC-seq (and Multiome ATAC) libraries, a minimum of 25,000 reads/nucleus was targeted. Multiome GEX libraries were targeted to a minimum 20,000 reads/nucleus. MULTI-ATAC and MULTI-seq barcode libraries were each sequenced to a target depth of at least 5,000 reads/nucleus.

FASTQs from the Multiome pilot experiment were aligned with Cell Ranger ARC (v2.0.1) to a mm10 reference assembly modified as described previously^74^ to properly align mitochondrial reads. FASTQs from all other experiments were aligned with Cell Ranger ATAC (v2.0.0, v2.1.0) or Cell Ranger ARC (v2.0.1) to the refdata-cellranger-arc-GRCh38-2020-A-2.0.0 reference assembly provided by 10x Genomics.

FASTQs from MULTI-ATAC and MULTI-seq barcode libraries were processed, aligned, and quality-controlled using deMULTIplex2^26^ before downstream sample-demultiplexing using the same software.

### scATAC-seq analysis pipeline

All scATAC-seq experiments were processed through a similar analytical pipeline before performing ad hoc analyses pertaining to each experimental design. In brief, each fragment file output by Cell Ranger ATAC or Cell Ranger ARC was processed with ArchR^21^ to produce an Arrow file containing a TileMatrix and GeneScoreMatrix. Single or multiple Arrow files from the same experiment were accessed and manipulated through an ArchRProject, allowing quality-control filtering based on per-cell metrics like TSS enrichment and fragment counts. Iterative Latent Semantic Indexing (iLSI) was used to produce a dimensionality reduction from the TileMatrix, and then typically dimensions 2-30 were used to generate a UMAP embedding for visualization purposes. The cell barcodes that passed QC were then fed into deMULTIplex2 and classified to their sample of origin utilizing the barcode counts tabulated from MULTI-ATAC reads. deMULTIplex2 classifications were then integrated into the ArchR project, and the project was subset to keep only the high-quality singlets identified from the MULTI-ATAC data before repeating iLSI and UMAP embedding. Downstream analyses typically included peak-calling via MACS2, motif deviation scoring via ChromVAR, and cell type annotation via marker analysis.

### Re-analysis of published datasets

For each of the 12 published datasets re-analyzed in this study, available pre-processed scATAC-seq data and metadata were downloaded from online repositories or as supplemental attachments in the form of fragment files, Seurat objects, or various per-cell or per-sample spreadsheets. When transposition batch information was not directly annotated, it was deduced based on the methods, computational tools, metadata, and experimental design information provided by authors in the accompanying publication and published analysis code.

When fragment files were readily available, datasets were processed with the standard ArchR pipeline (iLSI, clustering, and UMAP embedding), and were filtered to either only include high quality singlets, or only include cell barcodes identified by authors in supplementary files.

### PBMC donor genotypic demultiplexing

A list of cell barcodes and a BAM file containing position-sorted read alignments were fed into cellsnp-lite to genotype each cell based on a master list of 36.6M SNPs from the 1000 Genomes project (minMAF = 0.1, minCOUNT = 20). The resulting VCF file contained the variants detected in each cell and was processed with Vireo to probabilistically determine the donor identity of each cell, or assign it as a doublet.

### Drug/activation scoring

Because drugs in the Multiome drug screen were administered to PBMCs in the presence of immunostimulatory antibodies, we sought to isolate and quantitatively compare the effect of each drug dose on relative activation and all other drug-induced changes separately. To calculate the relative activation score, the accessibility of activation-associated marker genes for each cell type is aggregated by cell type and drug dose replicate. The mean aggregate value for resting control/DMSO(-) cells is then subtracted and then scores are normalized to the stimulated control/DMSO(+) cells. Thus, all drugs are scored by the same cell type-specific marker set and relative activation state can be compared. For the orthogonal drug score, we wanted to be able to compare paired inhibitors and PROTACs targeting the same enzyme. To do so, we selected the union marker set of each drug pair per cell type and excluded any markers that were involved in calculating the relative activation score. We then separately calculated the log2 fold-change in accessibility of the up- and down-regulated markers in this set relative to stimulated control/DMSO(+) cells. The absolute values of these two “up” and “down” drug scores were combined into a weighted average according to the relative proportion of up- or down-regulated markers in the set. The values plotted in Fig. 4A represent the average drug and activation scores for all 4 replicates per drug dose.

### Statistical analysis and data visualization

Statistical analysis and data visualization were performed in R (v.4.3.3). Single-cell chromatin accessibility and gene expression analysis across all experiments utilized the R packages ArchR^21^, Seurat^75^, and Signac^22^. Statistical tests and p-values are indicated in the text, figures, and figure legends.

## Supporting information

MULTI-ATAC Protocol

## Data and code availability

All analysis code and R objects necessary to reproduce key figures will be made available at github.com/Gartner-Lab/MULTI-ATAC. Processed sequencing data files will be uploaded to the Gene Expression Omnibus (GEO), and raw sequencing reads will be made available through the Short Read Archive (SRA).

## Acknowledgements

We thank Brittany Moser for supplying Jurkat and K562 cells used for testing the method and UCSF LARC for help with mouse husbandry. Sequencing was performed at the UCSF Center for Advanced Technology, and we appreciate their team for technical support and access to computational resources that aided in data processing and alignment. This work was funded by grants from the NIH (R01-GM135462-02 & R33-CA247744-01 to Z.J.G.), UCSF Center for Cellular Construction (no. DBI-1548297), Chan Zuckerberg Initiative, and UCSF Program for Breakthrough Biomedical Research (2021-2022 Postdoc Independent Research Grant to E.K.). Z.J.G. is a Chan Zuckerberg BioHub Investigator. We are grateful to all Gartner Lab members, as well as Danica Fujimori, Ryan Corces, and Vijay Ramani, for helpful feedback during the development of this work.

## Author contributions

D.N.C., C.S.M, and Z.J.G. conceived the project and designed the barcode and protocol. D.N.C. and E.K. performed the Multiome pilot experiment with hepatocytes. D.N.C., K.T.P., and Q.Z. performed the Multiome drug screen. D.N.C. and Z.J.G. designed all experiments and interpreted results. D.N.C. performed all other experiments and all data analysis. E.D.C. provided expertise on barcode design and next-generation sequencing. Z.J.G. and E.K. provided funding for experiments. D.N.C. and Z.J.G. wrote the manuscript with input from all authors.

---

**Supplementary Figure 1.**
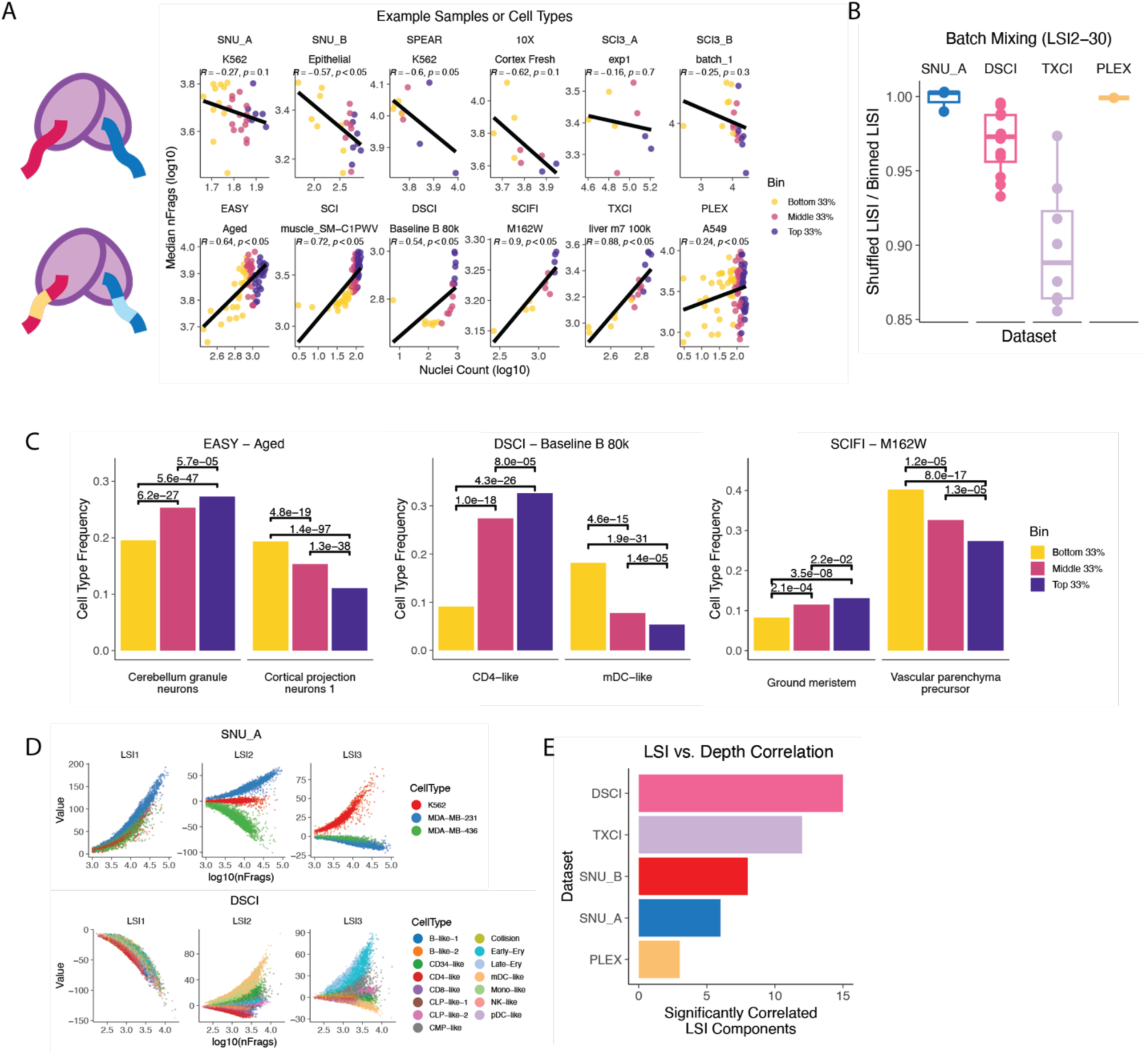
Batch effects linked to transposition batch size in published datasets. A. Example samples from each dataset. Points represent the nuclei count and median fragment count per transposition reaction, and are colored by transposition batch size tercile. Correlation coefficients and p-values from two-sided Pearson’s test. B. Batch mixing analysis, as in Fig. 1E), but excluding the 1^st^ LSI dimension as is standard practice due to correlation with depth. C. Demonstrative cell types from 3 other samples & datasets as in Fig. 1F), showing statistically significant changes in cell type frequency according to transposition batch tercile. P-values represent results from two-sided Chi-squared proportion tests. D. The 1^st^ LSI dimension obviously correlates with fragment count irrespective of cell type, whereas other dimensions show strong linear relationships with fragment count when separated by cell type. E. When aggregated by cell type, many LSI dimensions across 5 datasets correlate significantly with fragment count (R > 0.5, p < 0.05).

**Supplementary Figure 2.**
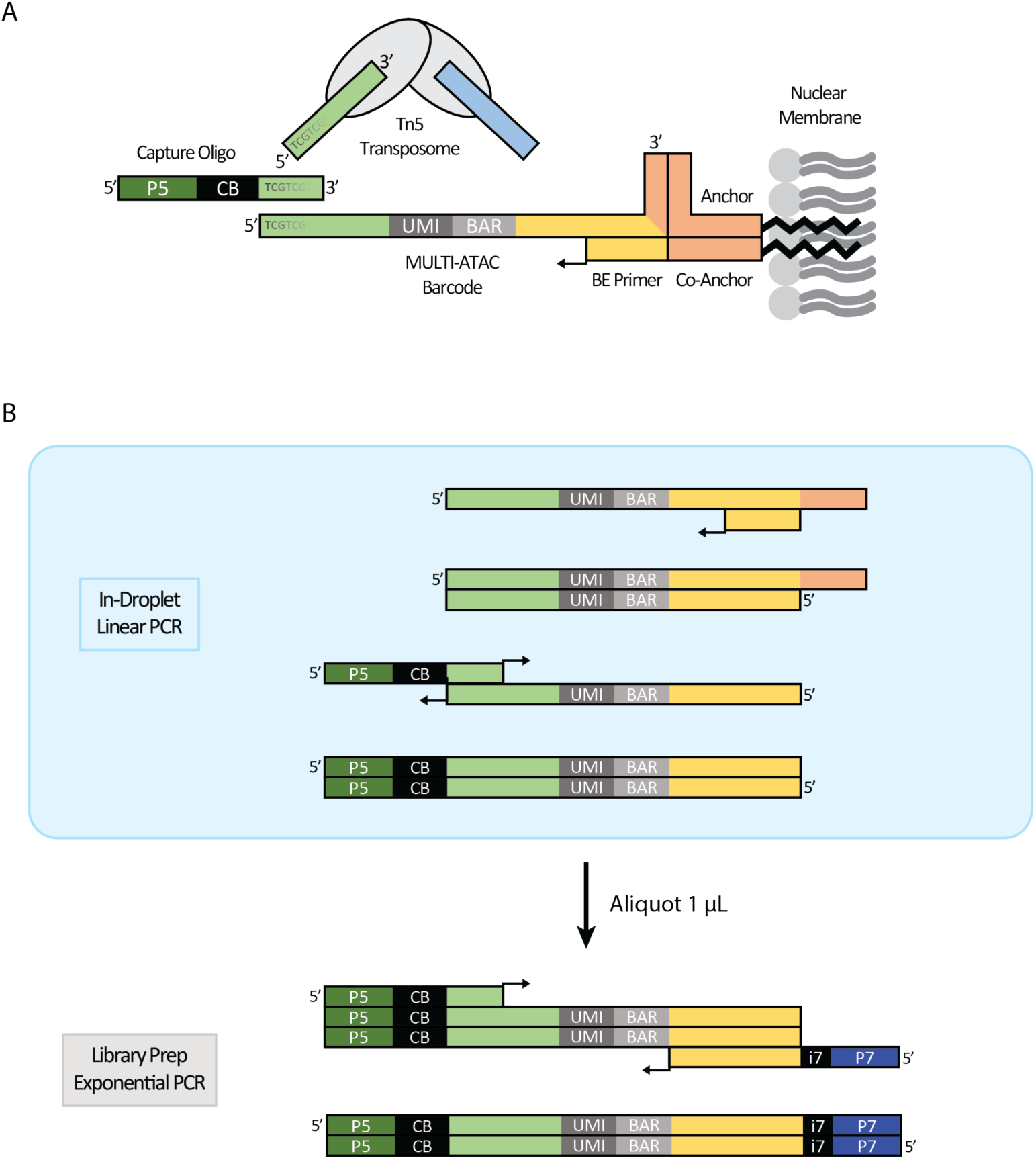
MULTI-ATAC Method Design. A. MULTI-ATAC barcodes are pre-hybridized to LMO Anchor and BE Primer oligos, and the full complex incorporates into nuclear membranes through step-wise addition with LMO Co-Anchor as in MULTI-seq. The orientation of the barcode prevents direct hybridization to the adapter oligos loaded into the Tn5 transposome. B. The pre-hybridized BE Primer is extended during in-droplet linear PCR to produce the complement strand required for priming with 10x Genomics capture oligos during subsequent rounds of linear PCR. After GEM incubation and cleanup, 1 *µ*L of library is aliquoted from the standard library preparation procedure to perform a separate PCR reaction with MULTI-ATAC specific sample-indexing primers.

**Supplementary Figure 3.**
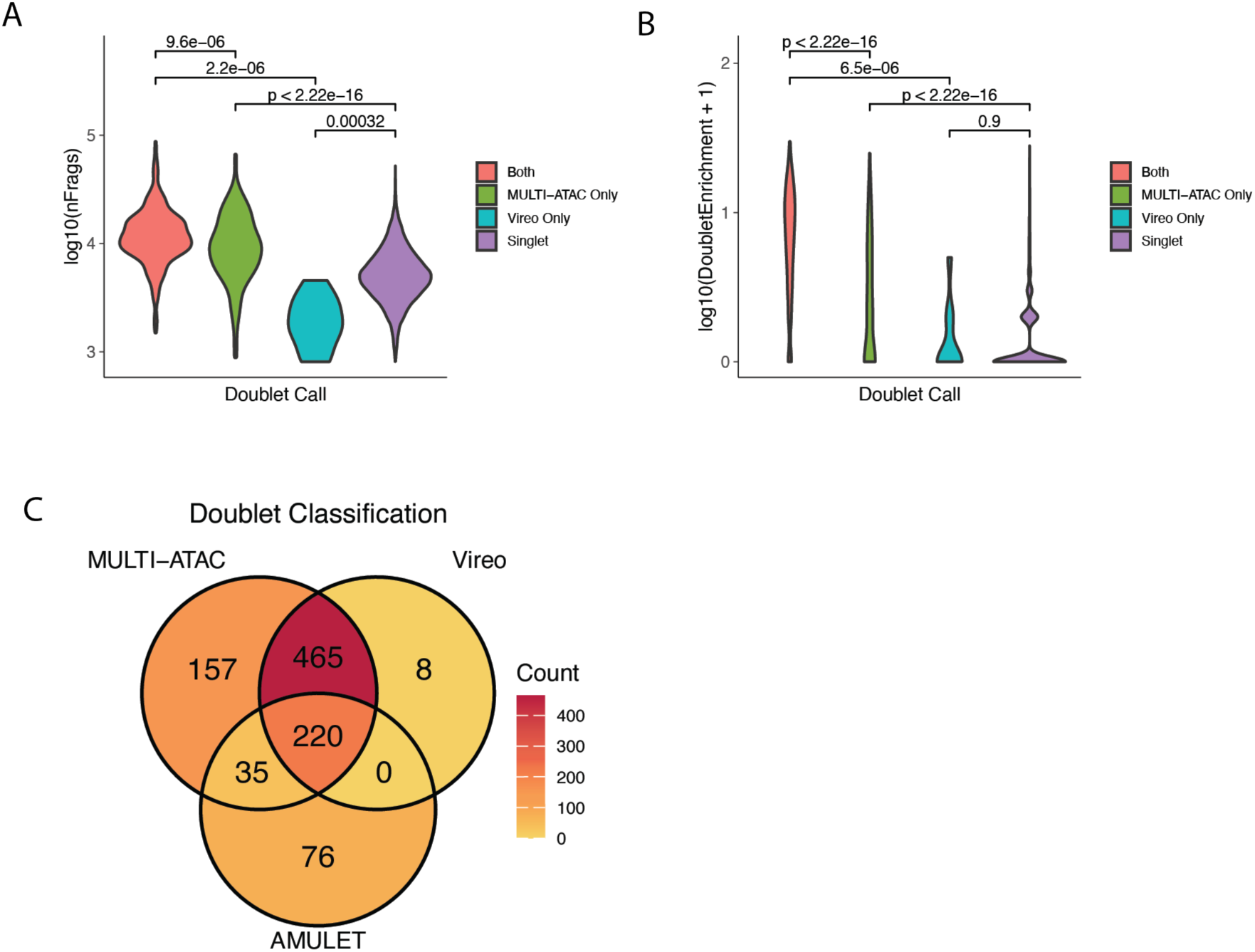
MULTI-ATAC identifies doublets not identified through fragment-based methods. A. Comparison of fragment counts for doublets classified by both MULTI-ATAC and Vireo, only MULTI-ATAC, only Vireo, or neither. Student’s t test. B. Comparison of DoubletEnrichment scores for doublets classified by both MULTI-ATAC and Vireo, only MULTI-ATAC, only Vireo, or neither. Student’s t test. C. Venn diagram comparing doublet classifications between MULTI-ATAC, Vireo, and AMULET. Notably there are no doublets agreed upon by Vireo and AMULET that MULTI-ATAC did not call.

**Supplementary Figure 4.**
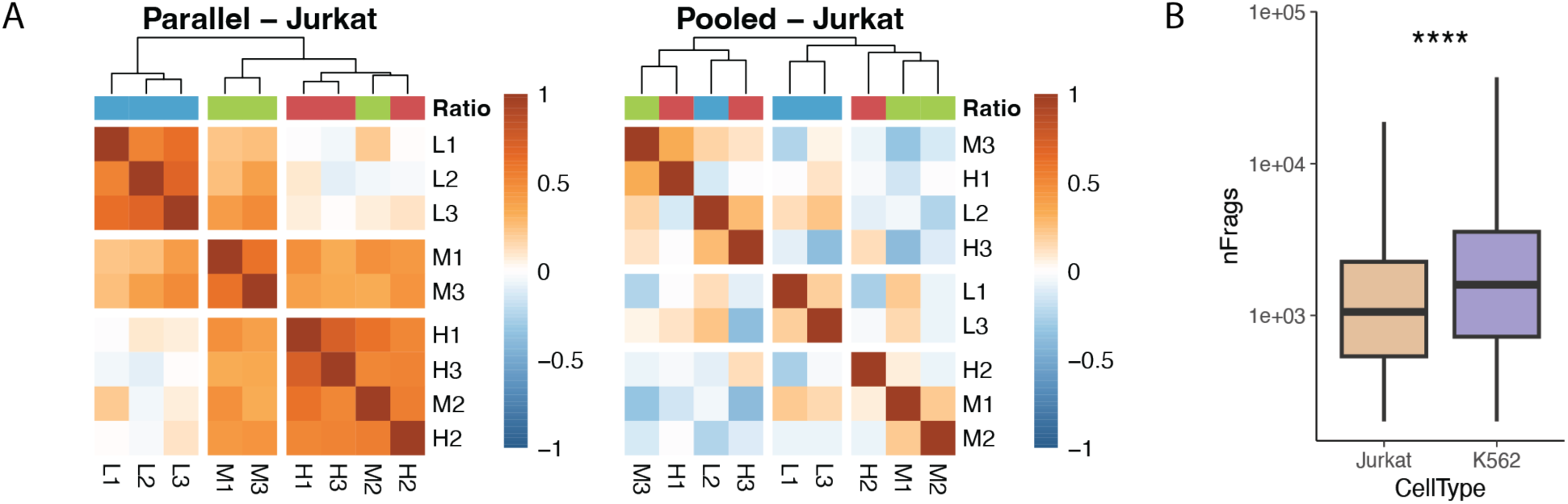
Pooled transposition eliminates batch effects. A. As in Fig. 2E), Jurkats cluster according to sample size in the Parallel library but not in the Pooled library. B. Jurkat nuclei yielded on average 36% fewer fragments than K562 nuclei, possibly making them more sensitive to quality control filtering.

**Supplementary Figure 5.**
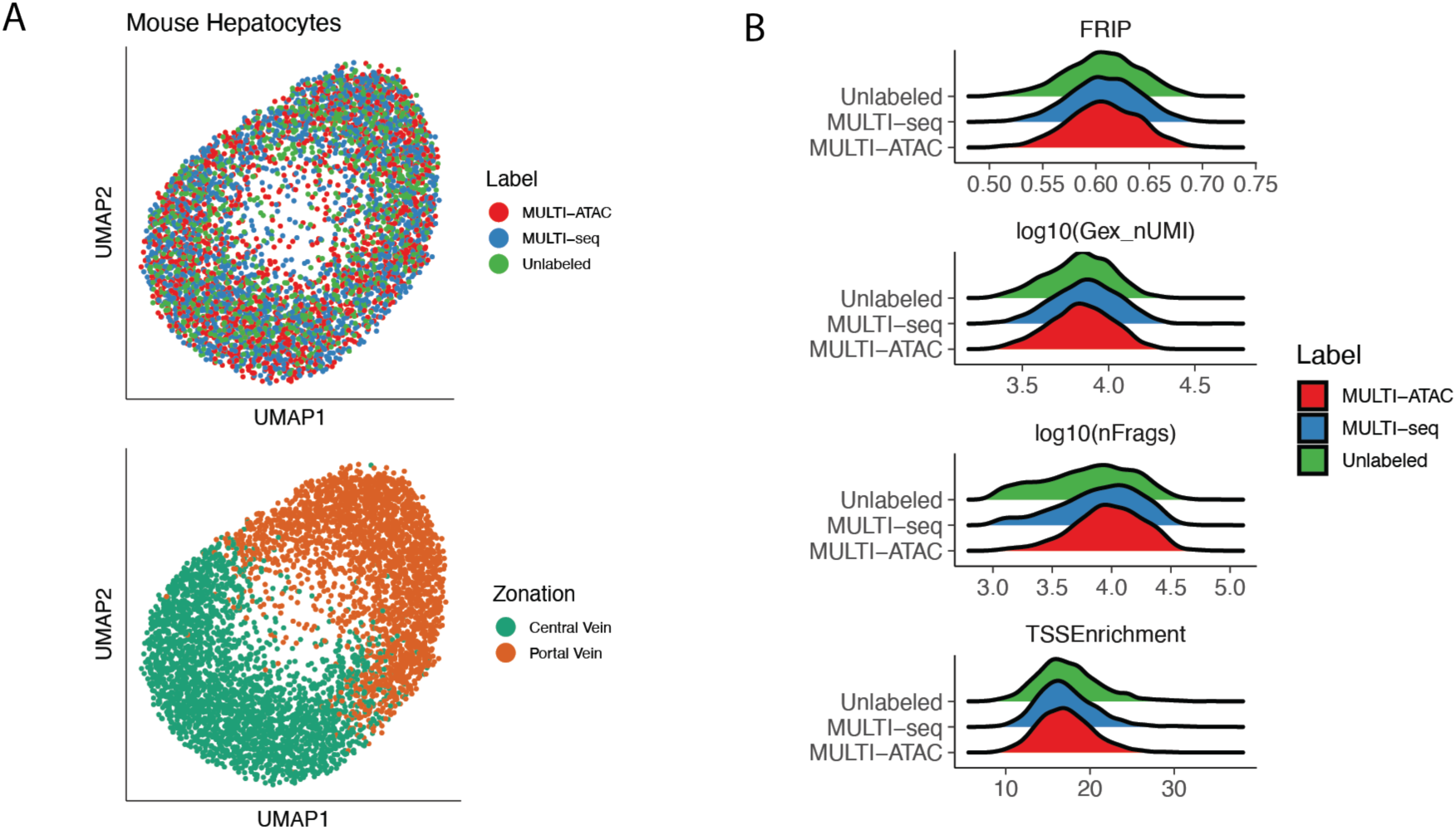
Multiome pilot experiment demonstrates similar performance to MULTI-seq. A. UMAP embeddings of GEX library captured in Multiome experiment shows separation of mouse hepatocytes by zonation markers (bottom), and homogenous mixing of cells labeled with either MULTI-ATAC barcodes, MULTI-seq barcodes, or neither. B. Comparison of ATAC and GEX library quality control metrics between hepatocytes labeled with MULTI-ATAC barcodes, MULTI-seq barcodes, or neither.

**Supplementary Figure 6.**
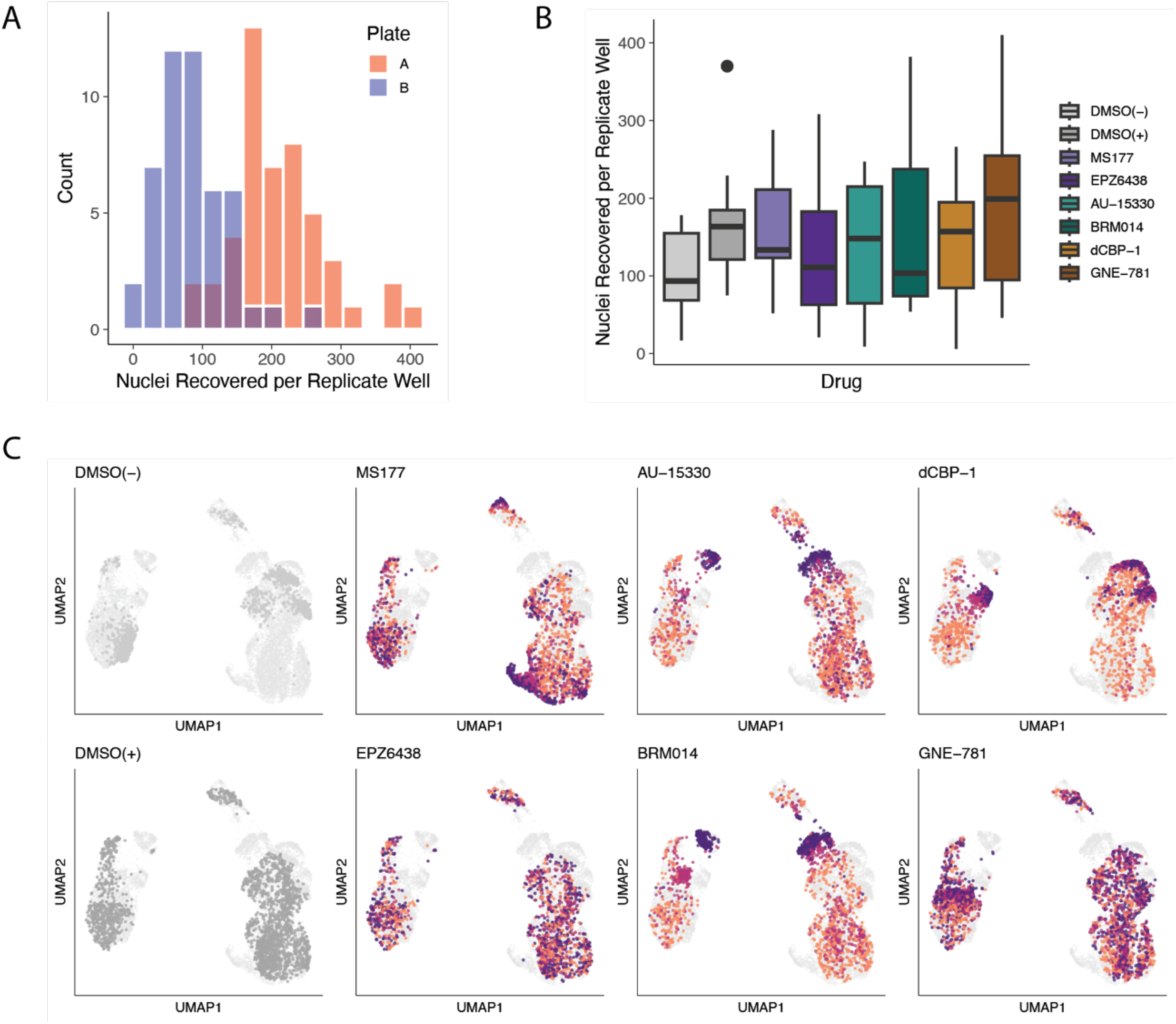
Recovery of replicates from each condition in final dataset. A. Overall, 148 ± 87 nuclei were recovered per replicate well with no dropouts. B. Overview of nuclei recovered per replicate well of each drug. C. UMAP embeddings for each drug and controls showing dose-dependent shifts in epigenetic state.

**Supplementary Figure 7.**
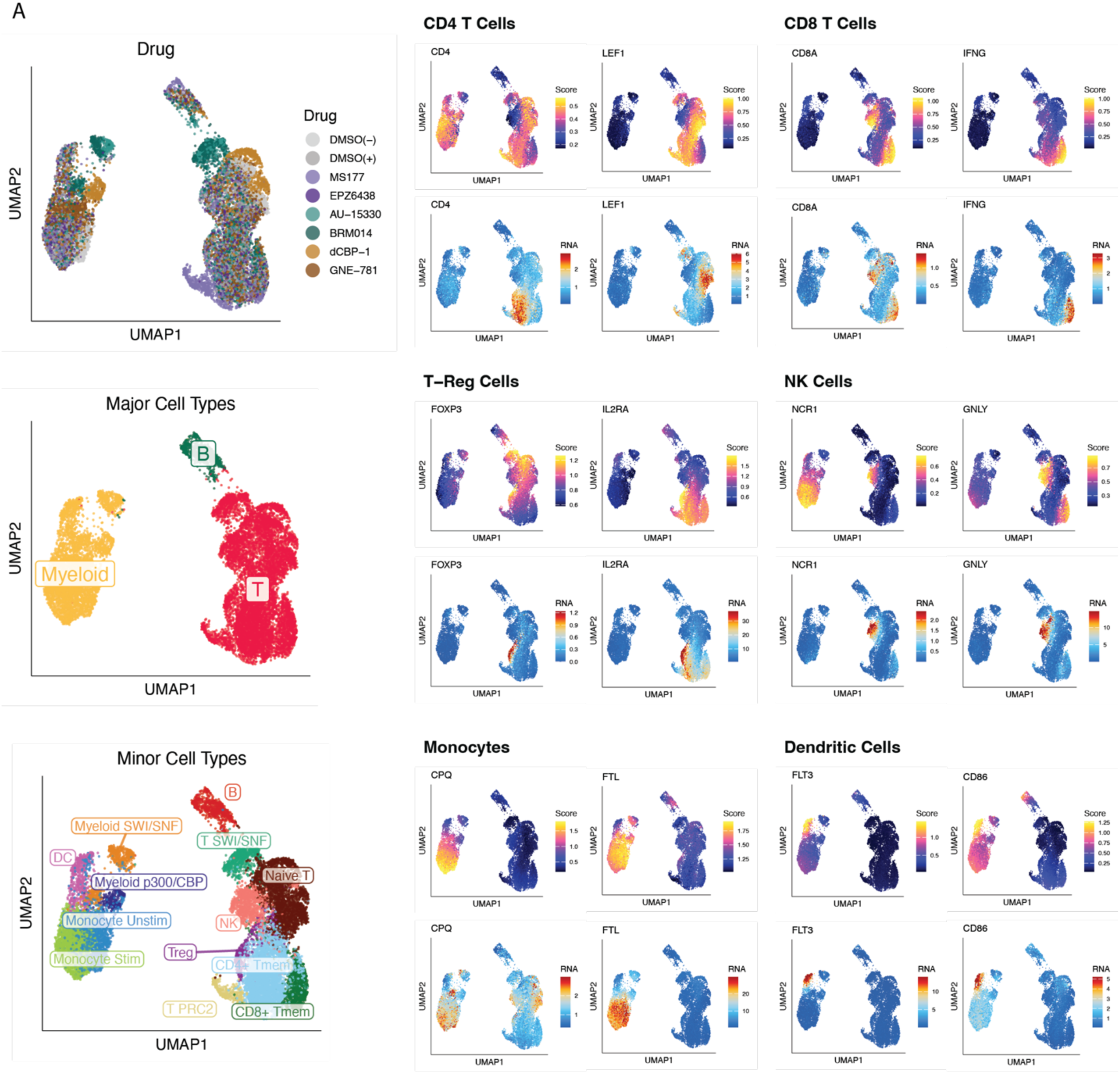
Major and minor cell type annotation using known markers. A. Canonical markers were assessed in terms of chromatin accessibility scores and RNA expression and used to annotate clusters as B cells, T cells (CD4+, CD8+, NK, and Treg), and Myeloid cells (Monocyte, DC). Several of the higher drug doses pushed cells into states that couldn’t be traced back to subtypes, and were annotated as such. For T and Myeloid populations, cells that clustered with resting control/DMSO(-) cells were annotated as Naïve/Unstimulated.

**Supplementary Figure 8.**
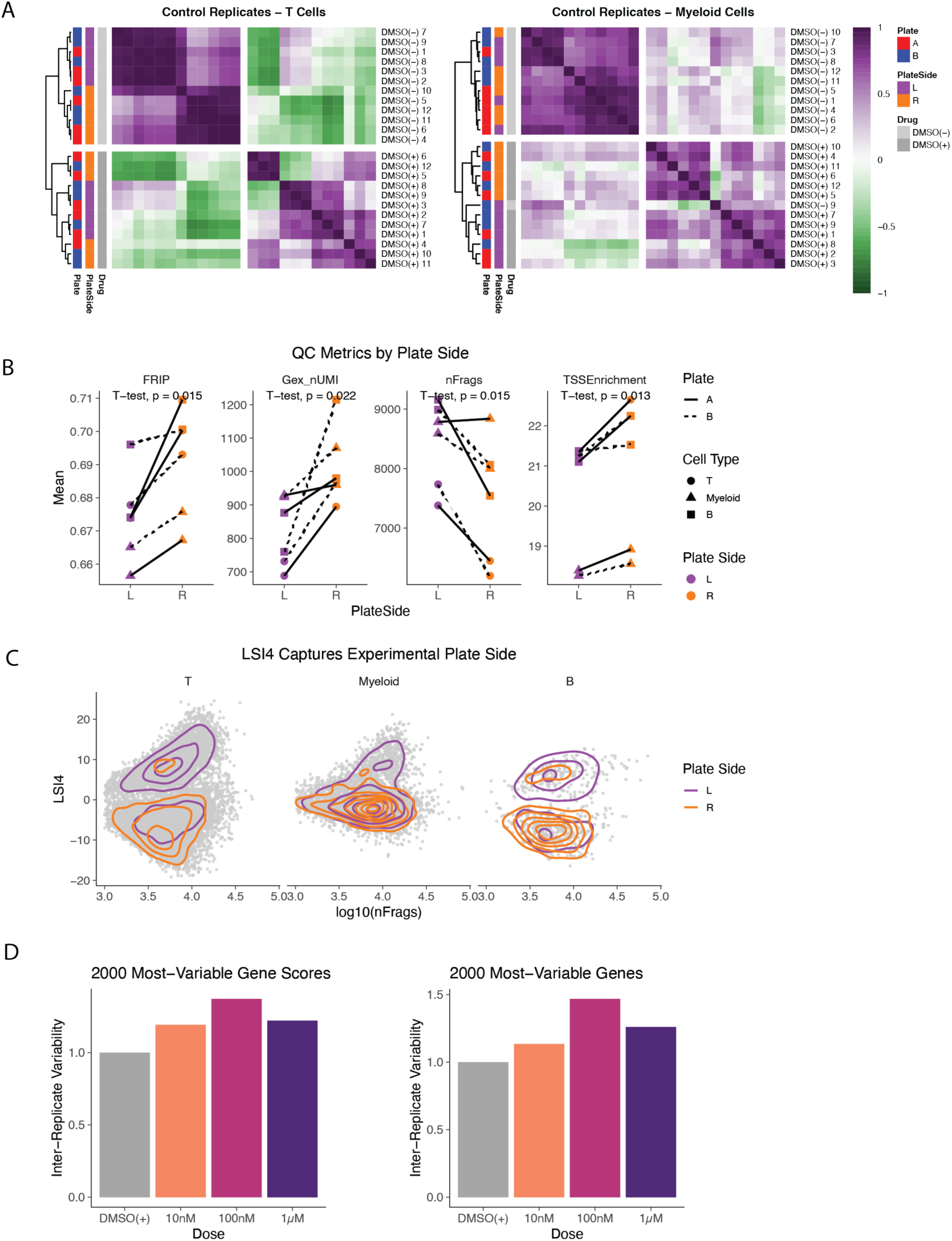
Inclusion of multiple replicates enables robust experimentation and statistical analysis. A. Control replicates were clustered by the correlation of their centroids in the LSI dimensionality reduction (as in Fig. 2E), and were found to cluster according to the sides of the plates (left vs right) they derived form. The mechanism behind this effect is not clear but could be linked to variable lysis or culture conditions. B. Cells from the left and right side of each plate differed significantly across various quality control metrics. C. Plate side seemed to be captured predominantly in LSI4, so this component was excluded from subsequent steps. This did not impact any downstream marker analyses, only visualization via UMAP and cell subtype annotation via clustering. D. Drug-dosed cells exhibited greater inter-replicate variability in gene accessibility and expression relative to control cells.

**Supplementary Figure 9.**
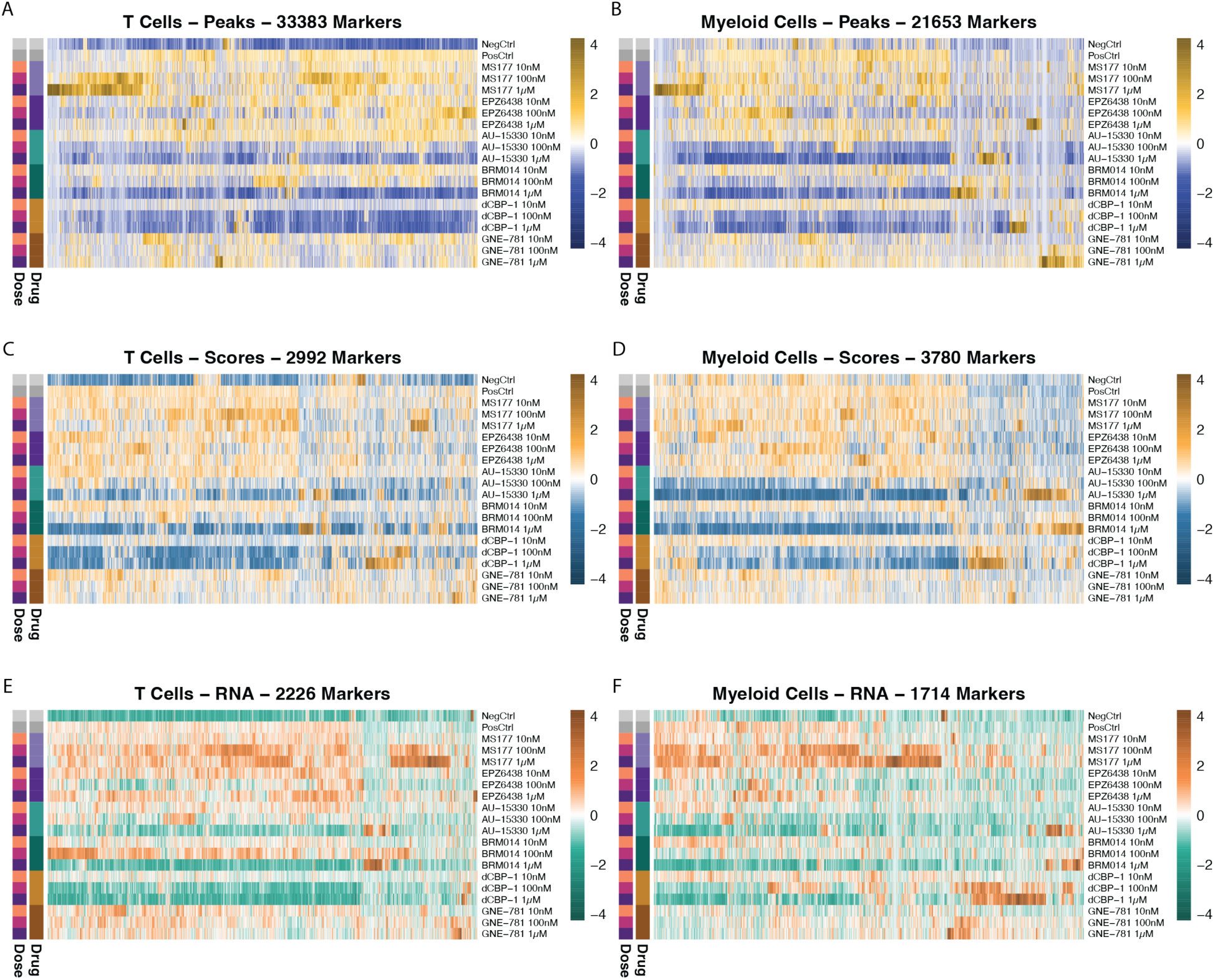
Heatmaps of markers significantly altered by immune activation and/or drug treatment. Heatmaps show Z-scaled median accessibility or expression values across replicates for each condition. A) 33,383 differentially accessible marker peaks in T cells (p < 0.01, Log2FC > 1) B) 21,653 differentially accessible marker peaks in Myeloid cells (p < 0.01, Log2FC > 1) C) 2,992 differentially accessible marker genes in T cells (p < 0.01, Log2FC > 1) D) 3,780 differentially accessible marker genes in Myeloid cells (p < 0.01, Log2FC > 1) E) 2,226 differentially expressed marker genes in T cells (p < 0.01, Log2FC > 1) F) 1,714 differentially expressed marker genes in Myeloid cells (p < 0.01, Log2FC > 1)

**Supplementary Figure 10.**
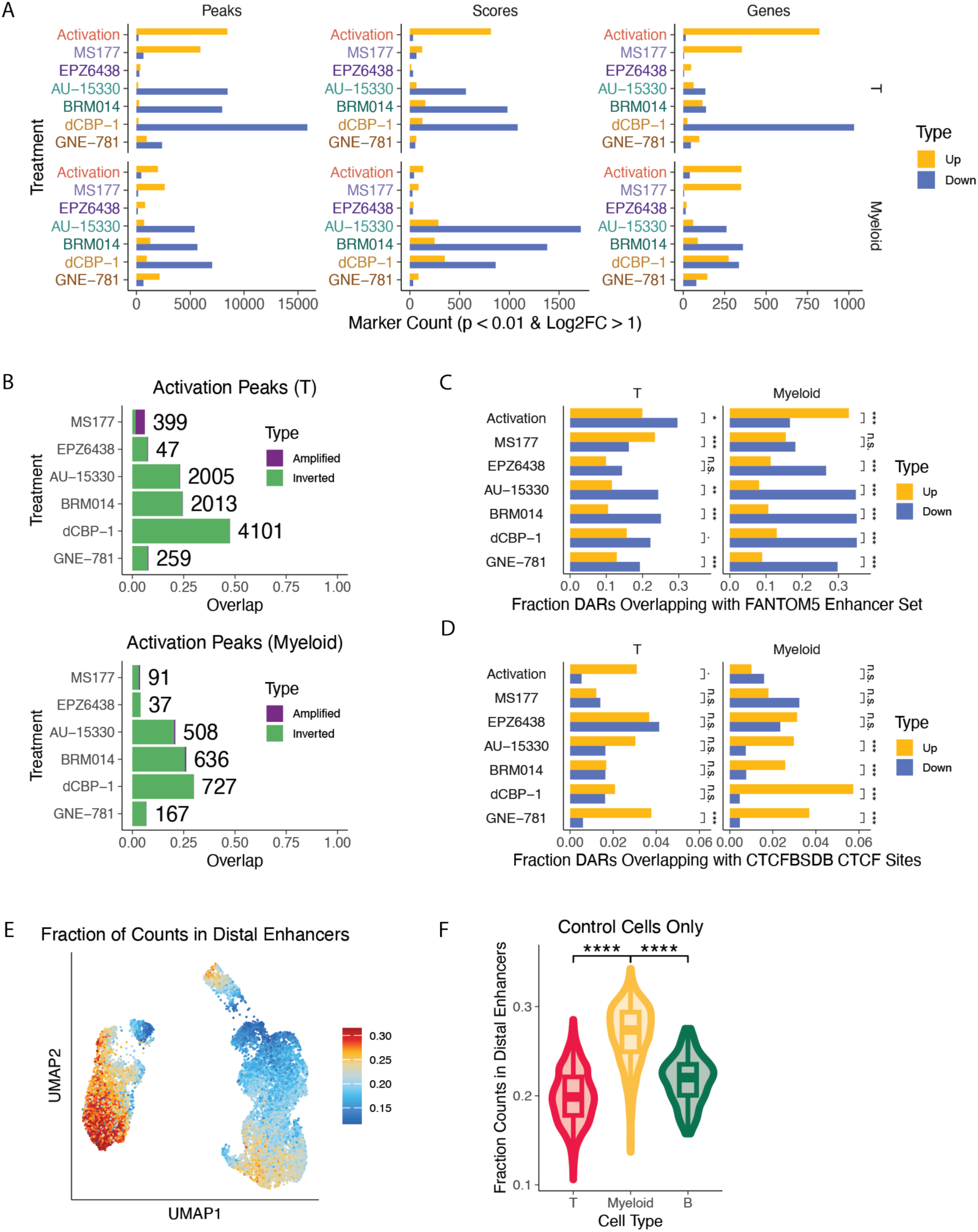
Characterization of marker features. A) Activation and MS177 treatment predominantly increased chromatin accessibility and gene expression, whereas treatment with drugs such as AU-15330, BRM014, and dCBP-1 largely had the opposite effect. B) A large portion of marker peaks in AU-15330, BRM014, and dCBP-1 reflect inversions of activation-associated chromatin accessibility changes. MS177 uniquely seems to further increase the accessibility of peaks already associated with T cell activation. C) Overlap of up- and downregulated peaks with FANTOM5 enhancer set. P-values represent results from two-sided Chi-squared proportion tests. D) Overlap of up- and downregulated peaks with CTCFSDB CTCF binding site database. P-values represent results from two-sided Chi-squared proportion tests. E) UMAP embedding of the per-cell fraction of fragments that overlap with distal enhancers from the CCRE database. F) Non-drugged myeloid cells exhibit a greater fraction of fragments coming from distal enhancers relative to T and B cells. Student’s t test.

**Supplementary Figure 11.**
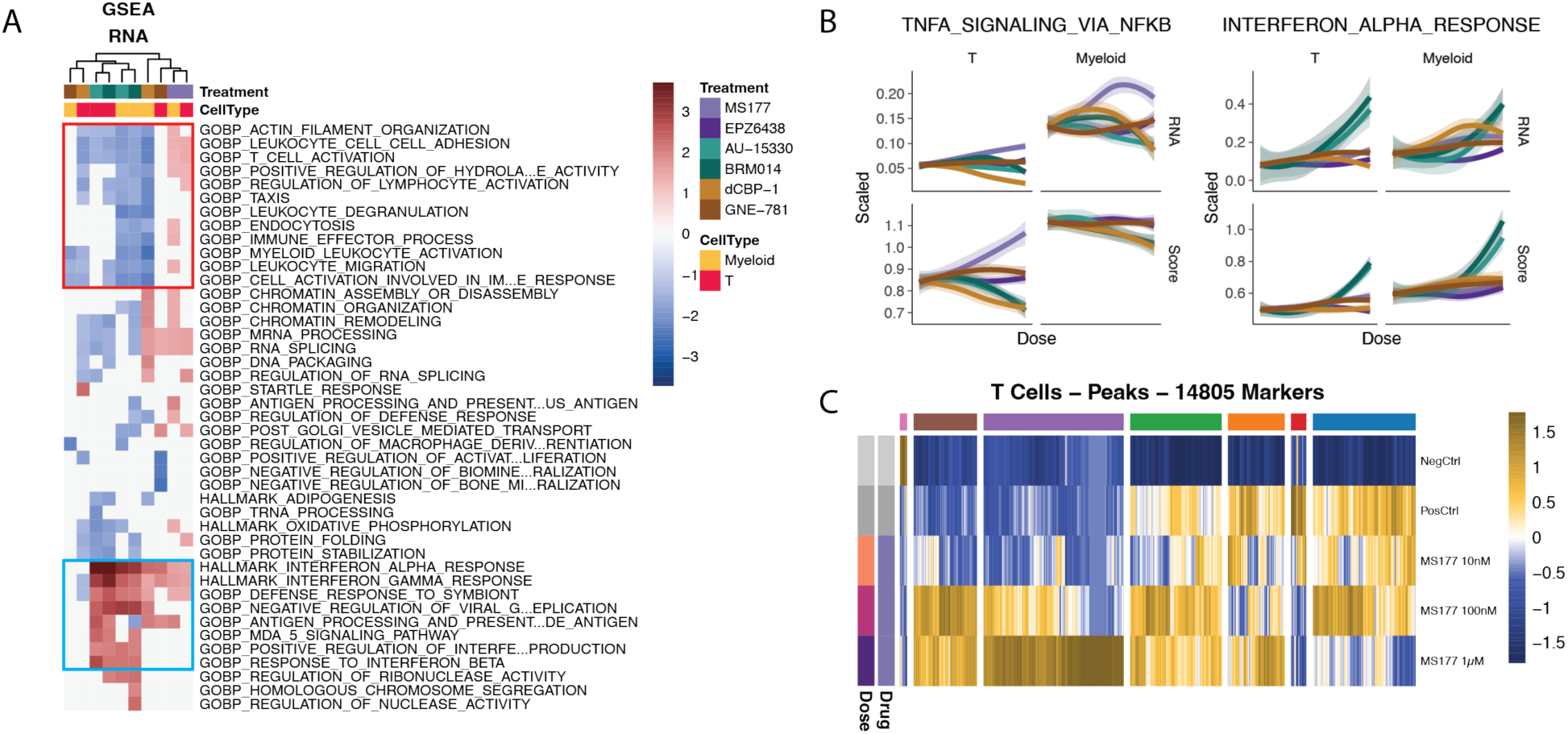
Pathway analysis of marker genes. A) As in Fig. 4B), gene set enrichment analysis (GSEA) for each drug and cell type of expressed marker genes ordered by statistical significance and negative vs positive slope. Statistically significant terms (p.adj. < 0.01) are colored by normalized enrichment score (NES). Red box – gene sets involved in immune cell activation; blue box – gene sets involved in type I interferon response. B) Similar to Fig. 4C), gene set expression across increasing drug dose; expression or accessibility of each gene at each drug dose was scaled relative to the activated controls, and then plotted as a function of dose. Trendlines plotted per drug via LOESS smoothing with span = 1.5. C) MS177- and activation-responsive marker peaks in T cells were hierarchically clustered into 7 groups for downstream motif enrichment analysis. Heatmap shows Z-scaled median accessibility values across replicates for each condition.

**Supplementary Figure 12.**
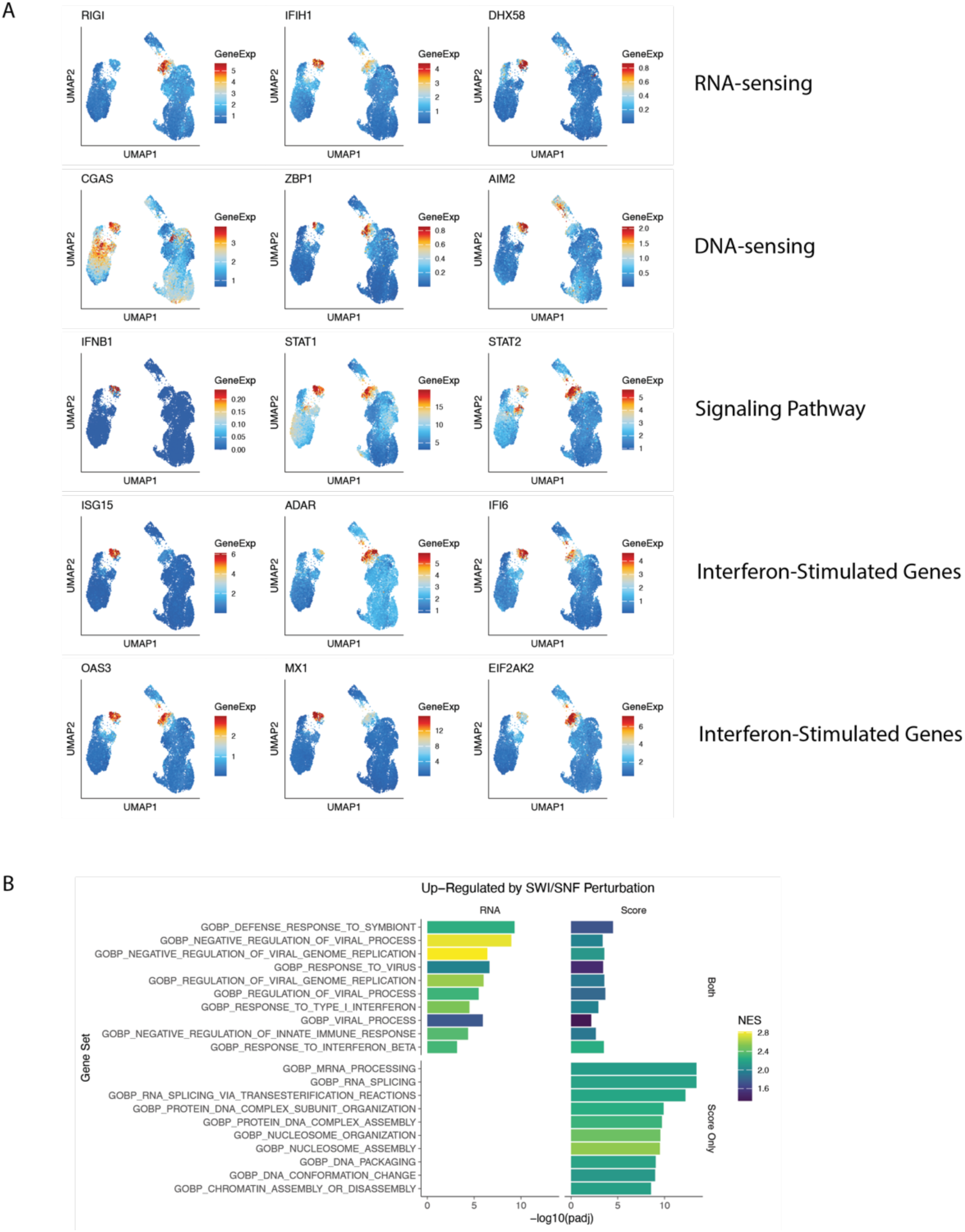
SWI/SNF perturbation upregulates genes and pathways related to Type I Interferon response. A. UMAP embeddings showing imputed RNA counts for 15 genes involved in Type I Interferon signal transduction and response. B. Gene sets determined to be upregulated by SWI/SNF perturbation through GSEA of RNA and Gene Score linear regression markers. Type I Interferon gene sets are upregulated in both modalities, whereas gene sets related to chromatin organization and RNA processing are only upregulated in accessibility but not expression.

**Supplementary Figure S13.**
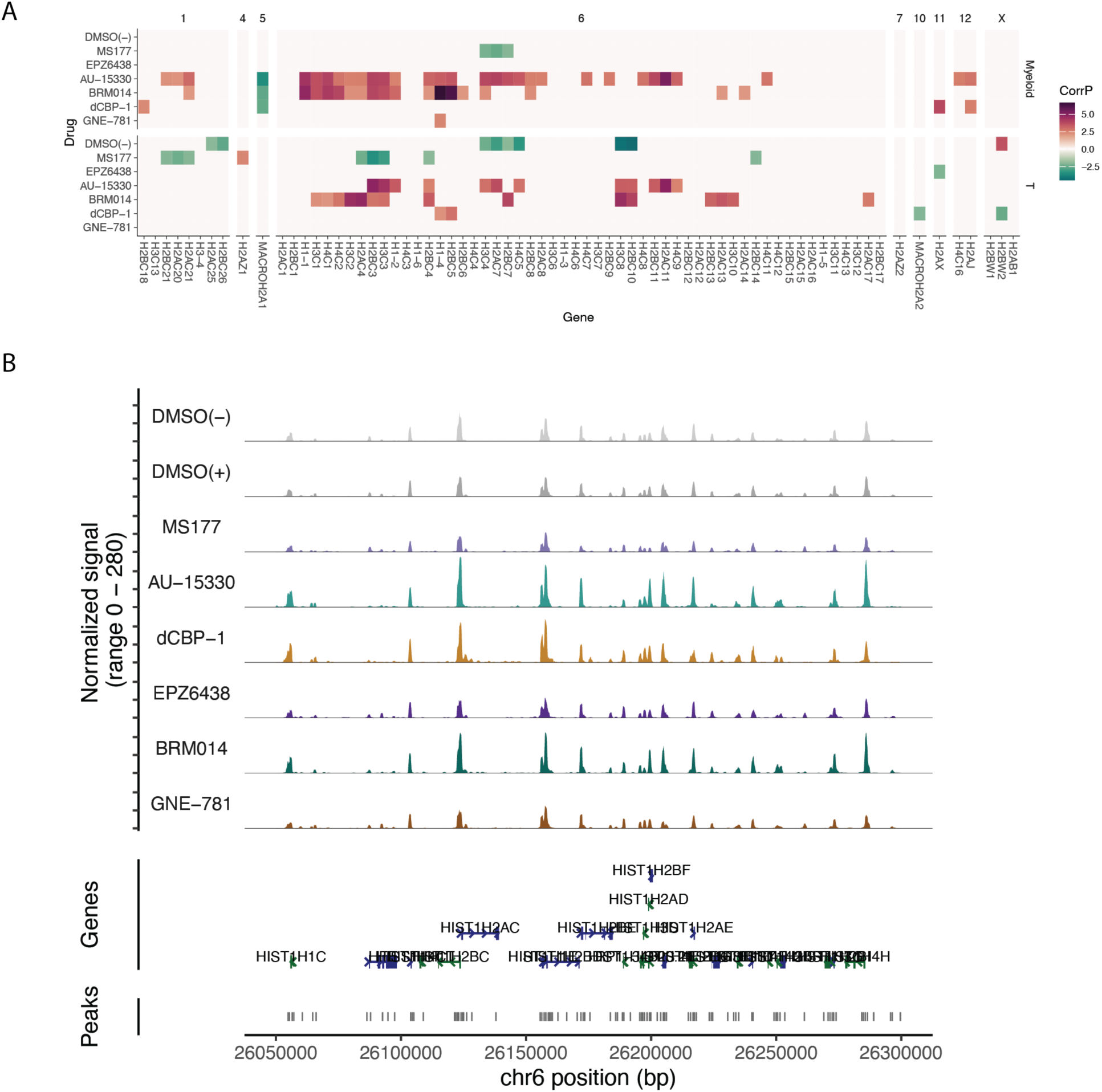
Replication-dependent histones among genes that gain accessibility from SWI/SNF perturbation. A. All histone genes – ordered by genomic location – colored by the direction and significance of their response in gene score/accessibility to increasing drug dose. B. Coverage plot of the HIST1 locus on chromosome 6 where most replication-dependent histone genes are located shows significant increases in accessibility, particularly for AU-15330 and BRM014 but also dCBP-1. Smoothing window for plotting = 1000 bp.

