## Supplementary material for "Reducing batch effects in single cell chromatin accessibility measurements by pooled transposition with MULTI-ATAC": MULTI-ATAC Protocol

Danny Conrad & Chris McGinnis
Gartner Lab, UCSF

*Updated 4/12/2024*

#### Introduction:

This protocol describes **MULTI-ATAC**, an adaptation of our MULTI-seq barcoding technology to enable pooling of samples for single-cell epigenomics. The method has been designed and validated for **10x Genomics scATAC-seq v1, v1.1, and v2** and **10x Genomics Multiome v1** kits. Much like MULTI-seq, barcodes integrate into cell and nuclear membranes via hybridization to lipid-modified oligonucleotides. The barcodes are captured and amplified alongside transposed chromatin fragments with only a few key modifications to the 10x kit protocols.

#### Method Notes:

##### *Pooled vs Parallel Transposition*

We provide two different workflows for multiplexing samples for scATAC-seq: pooled & parallel transposition. Sample barcoding enables nuclei from separate samples not only to be pooled when loading the 10x chip, but also to be transposed together in a single-pot reaction. This improves the workflow by:

- Using less Tn5 – an expensive/limiting reagent
- Reducing batch effects from running transpositions in parallel

However, due to the thermodynamic nature of the LMO interaction with nuclear membranes, the 37˚C incubation during transposition can lead to lower barcode UMI counts per nucleus. While we have had great success with the pooled method, the parallel transposition pipeline has marginally higher accuracy and is a viable alternative when the number of samples is low (i.e. 3-6 samples).

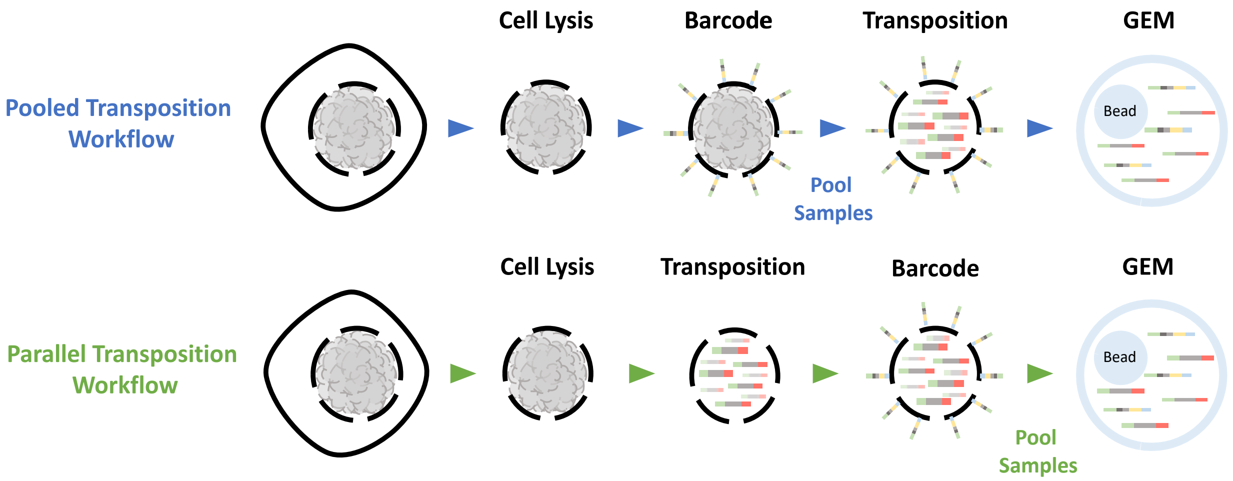


##### *Labeling Guidelines*

- Ensure nuclei are resuspended in BSA-free buffer with minimal clumping prior to labeling. LMOs partition quickly and clumping at this step can lead to heterogenous, noisy labeling. BSA binds long-chain fatty acids and is added back in after labeling to help clean up excess, unbound LMOs.
- Labeling volumes can be scaled up or down according to nuclei yields or experimental design, but we recommend keeping the final labeling concentration between 10-50nM. Additionally, try not to exceed target nuclei density during labeling (750-1000/μL) as this will dilute the labeling.
- Due to the magnitude of dilution required to prepare the Barcode Complex, we recommend pre-diluting an aliquot of your LMO Anchor, MULTI-ATAC Barcodes, and BE Primer stocks to the 1-10μM range. This ensures you can pipette realistic volumes ( >= 1 μL) to make just enough Barcode Complex for your experiment without wasting excess barcodes or LMOs. These adjusted "stock" concentrations are reflected in the example recipes provided for each workflow.

- As with MULTI-seq, we recommend testing labeling prior to commiting to a sequencing experiment, _*especially*_ when working with a new biological material or a custom nuclei isolation method. Prep nuclei and follow the labeling protocol but replace the barcode with a fluorophore-conjugated oligo (see "Sequences"). This allows a subjective assessment of labeling by flow cytometry. "Good" labeling shows strong signal over unlabeled controls and has a relatively unimodal, tight distribution.

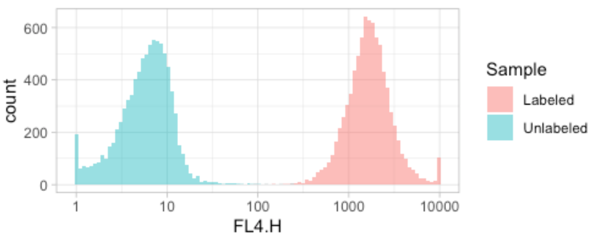


***Superloading***
As demonstrated with MULTI-seq, barcoding enables superloading by allowing high-confidence removal of most doublets. However, nuclei are inherently stickier than whole cells, and so increasing the density of nuclei loaded into each lane increases the occurance of multiplets and the risk of microfluidic lane clog. This will be highly sample- and nuclei-prep-dependent, so feel free to give it a try if so inclined. For reference, we think aiming to recover ~25,000 by loading ~40,000 adequately maximizes cell throughput without wasting too many sequencing reads on doublets (Kang *et al. Nat. Biotechnol.* **2018**, 36, 89). See also the Satija Lab website for calculating cost/cell.

***# Nuclei per Sample***
Per sample cell recovery will typically fall between 0.5X and 2X of your target. For example, if you want to get 500 cells per sample in a 96-sample experiment, you need to aim for 48,000 total cells. This should result in roughly 250-1000 cells recovered per sample. Consider the complexity of your sample when deciding on the number of cells per sample to target, starting with how many cells you hope to obtain from the lowest abundance cell type in your sample. For example, if you want an average of 50 cells from a cell type that makes up ~5% of your sample, then you would need to target 1000 cells per sample to obtain 25-100 cells per sample of your rare cell type. *SCOPIT is useful for this calculation.*

***Low Cell Input / Nuclei Yield***
When a sample has few total nuclei ($10^3$-$10^4$), our best advice is to scale down the labeling reaction, quench labeling by diluting samples with a large volume of 2% BSA in PBS, then immediately combine with other samples for pooled transposition.

***Sequencing***
Aim to sequence barcode libraries at ~5000 reads/nucleus. TruSeq-X primers (see below) incorporate library-specific i7 indices that allow barcode libraries from separate lanes to be pooled with each other and with corresponding scATAC-seq libraries for sequencing. Minimum cycle allocations: 16x16x16x6 (i5xR1xR2xi7)

***Data Processing / Demultiplexing***
We recently developed deMULTIplex2, a novel best-in-class demultiplexing algorithm for multiplexed single-cell sequencing methods (including MULTI-seq, Cell Hashing, etc.) with native support for MULTI-ATAC libraries. Implemented in R, deMULTIplex2 can generate sample classifications from raw FASTQs in a few minutes with just 3 steps.

- Read the publication: Zhu, et al. *Genome Biology* (2024)
- Install the package: GitHub

***Doublet Detection***
Heterotypic doublet/multiplet detection rate is directly proportional to the number of samples pooled. For example, doublets in an experiment with 12 sample barcodes have a 1 in 12 chance of having the same sample barcode. Therefore, the theoretical maximum detection rate is 11 in 12 or 91.7%. Tools such as AMULET & ArchR can complement MULTI-ATAC doublet detection, especially for homotypic (same-sample) doublets.

**<u>Reagents</u>**:

- 10x Genomics
  - Chromium Next GEM Single Cell ATAC (v1, v1.1, or v2) or Multiome ATAC + Gene Expression (v1)
  - Appropriate chips and index sets

- Library prep reagents
  - *Refer to appropriate 10x Genomics ATAC/Multiome User Guide*

- Nuclei isolation reagents
  - *Refer to appropriate 10x Genomics (or other) Nuclei Isolation Protocol*

- PCR strip tubes & 1.5mL Eppendorf tubes

- Kapa HiFi HotStart ReadyMix (2X)

- MULTI-seq Lipid-Modified Oligos (LMOs) (MilliporeSigma, cat#LMO001)

- MULTI-ATAC Barcodes & Accessory Oligos
  - order from IDT or other vendor with standard desalting
  - see "MULTI-ATAC_Sequences" spreadsheet for validated Barcode & TruSeq primer sequences
  - see below for Accessory Oligo sequences

- <u>Optional:</u> Illumina Tagment DNA TDE1 Enzyme & Buffer Kit (Illumina, cat# 20034197/20034198)
  - when excess Tn5 enzyme & buffer are needed to perform many **parallel** transposition reactions

### <u>Sequences</u>:

#### LMOs

Anchor LMO:

**Barcode Hybridization – Co-Anchor Hybridization – Lipid**
5′-TGGAATTCTCGGGTGCCAAGGgtaacgatccagctgtcact-{Lipid}-3′

CoAnchor LMO:

**Lipid – Anchor Hybridization**
5′-{Lipid}-AGTGACAGCTGGATCGTTAC-3′

#### Accessory Oligos

SI-PCR-B Primer

**Illumina P5**
5′-AATGATACGGCGACCACCGAGA-3′

TruSeq-[X] Primer

**Illumina P7 – i7 Index – TruSeq (Multiplexing PCR Primer 2.0)**
5′-CAAGCAGAAGACGGCATACGAGATXXXXXXGTGACTGGAGTTCAGACGTGTGCTCTTCCGATCT-3′

MULTI-ATAC Barcode Oligos
*Note: ‘N’ refers to random bases to allow barcode UMI counting, ‘X’ refers to predetermined bases for MULTI-ATAC Sample Barcodes*

***Gel Bead Oligo Capture*** **– Nextera Read 1N –** UMI **– SB – TruSeq (Multiplexing PCR Primer 2.0) – Anchor Hybridization (smallRNA R2)**
*5′-TCGTCGGCAGCGTC*AGATGTGTATAAGAGACAGNNNNNNNNXXXXXXXXAGATCGGAAGAGCACACGTCTGAACTCCAGTCACCCTTGGCACCCGAGAATTCCA-3′

Barcode Extension (BE) Primer

**Partial TruSeq (Multiplexing PCR Primer 2.0)**
5′-GTGACTGGAGTTCAGACGTGTGC-3′

Fluorophore-Conjugated Oligo
*Optional: to test labeling via flow cytometry*

5′-CCTTGGCACCCGAGAATTCCA-{FLUOROPHORE}-3′

(AlexaFluor 488 & 647 work in our hands)

#### Part 0: 10x Protocol Modifications

Once pooled, transposed nuclei have been loaded into the 10x chip and partitioned into GEMs, you more-or-less follow the protocol provided by 10x with two important exceptions:

- CRITICAL: In order to generate the matching MULTI-ATAC barcode library for each ATAC library you produce in a 10x run (**Part 5** of this protocol), a 1 μL aliquot needs to be taken prior to performing Sample Indexing PCR of the ATAC library. If you skip this step, it is unlikely your barcodes will be salvageable and the sample identity of your nuclei will be lost.

  When to take the aliquot:

  **scATAC-seq: Step 3.2o**
  Since the rest of the 40μL volume will be used up to generate the ATAC library, you can choose to run the Barcode Library Prep before completing the rest of ATAC Library Construction to make sure you successfully produced a MULTI-ATAC barcode library. Putting the ATAC Library Construction on hold gives you a second chance at barcode library prep if something were to go wrong.

  **Multiome: Step 4.2-4.3**
  This protocol is more forgiving due to the excess pre-amplified library that you get at the end of Step 4.3. 160 μL are produced but only 75 μL are used for downstream ATAC & GEX library construction, so you can draw repeat 1 μL aliquots from this excess if necessary. However, we've found a slightly better barcode library is produced by taking the 1 μL aliquot immediately after Pre-Amplification PCR, before the SPRI clean-up in Step 4.3. Both are viable options.

- When performing Sample Index PCR of the ATAC library, we prefer to replace the 10x-supplied SI-PCR Primer B (PN-2000128) with our own 100μM SI-PCR-B primer (see Accessory Oligos). The sequence and volume used are the same, but this increases the concentration of the primer in the reaction. We believe this prevents the rare occurrence of barcodes out-competing gDNA fragments for forward primers during Sample Index PCR, which inhibits exponential amplification.

#### Part 1: Nuclei Purification

*Note: This is adapted from 10x Genomics note #CG000169, "Nuclei Isolation for Single Cell ATAC Sequencing"; see this document for buffer recipes. Lysis time should be optimized to your sample. Other nuclei isolation methods can also be used but we advise users to validate LMO labeling (see "Method Notes" above).*

1. Wash cells once with chilled PBS
2. Aliquot 500k cells per sample into 1.5mL Eppendorf tubes
3. Pellet cells (4 min, 300g, 4˚C)
4. Remove supernatant
5. Resuspend in 100 μL chilled Lysis Buffer, pipette mix 10X
6. Incubate 3-5 min on ice
7. Add 1 mL chilled Wash Buffer to lysed cells, pipette mix 5X
8. Pellet nuclei (4 min, 500g, 4˚C)
9. Resuspend in chilled PBS, assuming ~50% nuclei loss
10. Count each sample
11. Adjust concentration of samples according to desired workflow:
    a. Pooled transposition: 750-1000 nuclei/μL
    b. Parallel transposition: see "Nuclei Concentration Guidelines" table in 10x scATAC-seq User Guide

### Pooled Transposition Workflow

#### Part 2: Barcoding

Prepare MULTI-ATAC barcoding reagents

- 400nM Barcode Complexes
  - Combine LMO Anchor, MULTI-ATAC Barcode, and BE Primer at 2:1:2 molar ratio
    - Maximizes complex formation with excess anchor and primer
  - One unique Barcode Complex per sample
- 800nM LMO CoAnchor

Example:

| | | [Stock] μM | Labeling Molar Ratio | [Labeling Solution] nM | Volume (μL) |
|---|---|---|---|---|---|
| Barcode Complex (10 μL per sample) | LMO Anchor | 10 | 2 | 800 | 2 |
| | MULTI-ATAC Barcode | 10 | 1 | 400 | 1 |
| | BE Primer | 10 | 2 | 800 | 2 |
| | Water or PBS | | | | 20 |
| | | | | Total | 25 |
| CoAnchor Solution (10 μL x nSamples) | LMO CoAnchor | 50 | 2 | 800 | 8 |
| | Water or PBS | | | | 492 |
| | | | | Total | 500 |

Barcoding

1. Aliquot 190 μL of each nuclei suspension into 1.5mL Eppendorf tubes
2. Add 10 μL 400nM Barcode Complex to nuclei for a final concentration of 20nM, immediately pipette mix or pulse vortex 10X
3. Incubate on ice, 5'
4. Add 10 μL 800nM CoAnchor Solution to nuclei, immediately pipette mix or pulse vortex 10X
5. Incubate on ice, 5'
6. Quench labeling by adding 1.2 mL 2% BSA in PBS
7. Pellet nuclei (4 min, 500g, 4˚C)
8. Aspirate supernatant, resuspend in 100-200 μL 2% BSA in PBS (count if desired)

#### Part 3: Pooling

9. Pool samples in one or more tubes at desired ratios based on experimental goals
10. Pellet pooled nuclei (4 min, 500g, 4˚C)
11. Resuspend in 1x Nuclei Buffer to 1/10-1/20 the original pooled volume
12. Count and adjust volume with 1x Nuclei Buffer such that 5 μL contains the number of nuclei you'd like to load per lane

#### Part 4: Transposition

13. Proceed immediately with "Step 1 – Transposition" of 10x Protocol, using 5 μL pooled nuclei

*NOTE: If transposing with 3$^{rd}$-party Tn5/buffer, resuspend transposed nuclei in a 1:2 solution of 1X Nuclei Buffer and ATAC Buffer B before loading 10x chip to ensure successful in-droplet linear PCR*

### Parallel Transposition Workflow

#### Part 2: Transposition

*Note: "Illumina" transposition is adapted from the Omni-ATAC protocol by Corces et al. (2017)*

> *NOTE: To use 10x-supplied transposition reagents, skip this section and proceed with 10x scATAC-seq Protocol "Step 1 - Transposition", then continue to Part 3: Barcoding*

1. Prepare Illumina Transposition Mix (13 $\mu$L per sample):
   - 7.5 $\mu$L 2X Tagment DNA Buffer
   - 2.95 $\mu$L 1X PBS
   - 0.15 $\mu$L 10% Tween-20
   - 0.15 $\mu$L 1% Digitonin
   - 0.75 $\mu$L Tagment DNA Enzyme 1
   - 1.5 $\mu$L Nuclease-Free $H_2O$
2. Add 13 $\mu$L transposition mix to 2 $\mu$L nuclei in fresh 1.5 mL Eppendorf, gently pipette mix 6X
3. Incubate at 37˚C for 1hr
4. Transfer transposed nuclei to ice

#### Part 3: Barcoding

Prepare MULTI-ATAC barcoding reagents

- 100nM Barcode Complexes
  - Combine LMO Anchor, MULTI-ATAC Barcode, and BE Primer at 2:1:2 molar ratio
    - Maximizes complex formation with excess anchor and primer
  - One unique Barcode Complex per sample
- 250nM LMO CoAnchor

Example:

| | | [Stock] $\mu$M | Final Molar Ratio | [Labeling Solution] nM | Volume ($\mu$L) |
|---|---|---|---|---|---|
| Barcode Complex (5 $\mu$L per sample) | LMO Anchor | 1 | 2 | 200 | 2 |
| | MULTI-ATAC Barcode | 1 | 1 | 100 | 1 |
| | BE Primer | 1 | 2 | 200 | 2 |
| | Water or PBS | | | | 5 |
| | | | | Total | 10 |
| CoAnchor Solution (5 $\mu$L x nSamples) | LMO CoAnchor | 50 | 2 | 250 | 1 |
| | Water or PBS | | | | 199 |
| | | | | Total | 200 |

Barcoding

5. Add 5 $\mu$L 100nM Barcode Complex to 15 $\mu$L transposed nuclei for a final concentration of 25nM, immediately pipette mix or pulse vortex 10X
6. Incubate on ice, 5'
7. Add 5 $\mu$L 250nM CoAnchor Solution to nuclei, immediately pipette mix or pulse vortex 10X
8. Incubate on ice, 5'
9. Quench labeling by adding 1.2 mL 2% BSA in PBS
10. Pellet nuclei (4 min, 500g, 4˚C)

11. Aspirate supernatant, resuspend in 100-200 $\mu$L 2% BSA in PBS

#### Part 4: Pooling

12. Pool samples in one or more tubes at desired ratios based on experimental goals
13. Pellet pooled nuclei (4 min, 500g, 4˚C)
14. *Carefully* aspirate as much supernatant as possible
15. Resuspend nuclei in ATAC Buffer B to a total volume of 30-50 $\mu$L*

*NOTE: Resuspending nuclei in ATAC Buffer B is necessary for successful in-droplet linear PCR*

16. Count and adjust volume with ATAC Buffer B such that 15 $\mu$L contains the number of nuclei you'd like to load per lane
17. Proceed immediately with "Step 2 - GEM Generation & Barcoding" of 10x Protocol, using 15 $\mu$L pooled transposed nuclei

#### Part 5: Barcode Library Prep

*CRITICAL: Follow 10x scATAC-seq Protocol through Step 3.2o (GEM Cleanup – SPRI selection), then STOP*

1. Remove 1 $\mu$L (2.5%) of scATAC-seq library for barcode library preparation

   *NOTE: Use remainder of ATAC-seq library for Step 4 of the 10x scATAC-seq Protocol*

2. Prepare following PCR reaction:

   - 2.5 $\mu$L 10$\mu$M SI-PCR-B
   - 2.5 $\mu$L 10$\mu$M TruSeq-X
   - 1 $\mu$L scATAC-seq library
   - 26.25 $\mu$L Kapa HiFi HotStart ReadyMix
   - 17.75 $\mu$L Nuclease-Free Water

3. Run MULTI-ATAC library prep PCR:

   1. 95˚C, 5:00
   2. 98˚C, 0:20
   3. 67˚C, 0:30
   4. 72˚C, 0:20
   5. Repeat Steps 2-4 x13
   6. 72˚C, 1:00
   7. 4˚C, Hold

4. Add 100 $\mu$L SPRI (for 2.0X clean-up), pipette mix 10x
5. Incubate at room temperature for 5'
6. Place on magnet (HIGH) until clear, remove supernatant
7. Add 200 $\mu$L 80% EtOH to beads, wait for 30"
8. Aspirate 80% EtOH, add another 200 $\mu$L 80% EtOH, wait for 30"
9. Aspirate 80% EtOH, centrifuge briefly, return to magnet (LOW)
10. Aspirate remaining 80% EtOH
11. Remove from magnet, resuspend beads in 20 $\mu$L Buffer EB
12. Incubate at room temperature for 2' to elute barcode library from beads
13. Return to magnet (LOW), transfer supernatant to fresh strip tubes

#### Part 6: Quantification & Sequencing

1. Make 1:5 dilution of barcode library in Buffer EB and analyze using Bioanalyzer or TapeStation
   a. Measure molarity/concentration of the peak around ~160-170 bp
2. Sequence barcode library at ~5000 reads/cell (see "Method Notes" for more details)

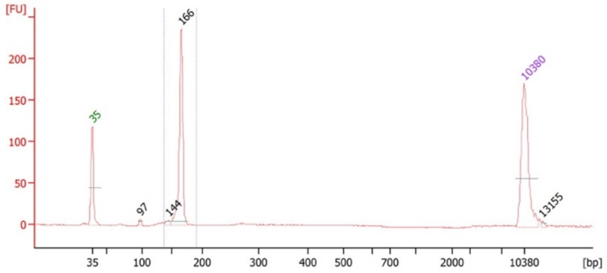


*Representative BA trace from MULTI-ATAC PBMC pilot experiment, from Conrad et al., 2024 (in preparation)*

### Appendix:

#### 96-well Barcoding (for Pooled Workflow)

Prepare MULTI-ATAC barcoding reagents

- 75nM Barcode Complexes
  - Combine LMO Anchor, MULTI-ATAC Barcode, and BE Primer at 2:1:2 molar ratio
    - Maximizes complex formation with excess anchor and primer
  - One unique Barcode Complex per sample
- 200nM LMO CoAnchor

Example:

| | | [Stock] μM | Labeling Molar Ratio | [Labeling Solution] nM | Volume (μL) |
|---|---|---|---|---|---|
| Barcode Complex (50 μL per sample) | LMO Anchor | 1 | 2 | 150 | 15 |
| | MULTI-ATAC Barcode | 1 | 1 | 75 | 7.5 |
| | BE Primer | 1 | 2 | 150 | 15 |
| | Water or PBS | | | | 62.5 |
| | | | | Total | 100 |
| CoAnchor Solution (50 μL x nSamples) | LMO CoAnchor | 50 | 2 | 800 | 8 |
| | Water or PBS | | | | 1992 |
| | | | | Total | 2000 |

Barcoding

14. Transfer 100 μL nuclei suspension to each well of a round-bottom 96-well plate
15. Add 50 μL 75nM Barcode Complex to nuclei for a final concentration of 25nM, gently pipette mix 5X
16. Incubate on ice, 5'
17. Add 50 μL 200nM CoAnchor Solution to nuclei, gently pipette mix 5X
18. Incubate on ice, 5'
19. Spin plate (4', 500g, 4˚C)
20. Carefully remove 195 μL of supernatant
21. Resuspend nuclei with 195 μL 2% BSA in PBS

Proceed with **Part 3: Pooling** & **Part 4: Transposition** of the **Pooled Transposition Workflow**
